# Chemically-induced Jamming and 3D printing of Granular Hydrogels: Microgels as Reservoirs of Volume

**DOI:** 10.64898/2026.09.28.754645

**Authors:** Micaela Fernandes, Julien Es Sayed, Armin Amirsadeghi, Rency Geevarghese, Joanna Żur-Pińska, Joop de Vries, Marcus Koch, Małgorzata Katarzyna Włodarczyk-Biegun

## Abstract

Granular hydrogels, made of jammed soft microparticles, are of great interest for 3D (bio)printing, as they combine ideal rheological properties and extensive modularity, yielding favorable microstructures for tissue engineering. Typically, the yield-stress properties of these materials, which facilitate printability, are defined by the preparation state and the initial particle content of the hydrogel. Here, we propose a granular hydrogel whose yield-stress and, consequently, printability and printed scaffolds’ shape retention can be controlled not only by the initial weight fraction of particles but also by their stimuli-responsive swelling. The responsive microgels are prepared from poly(*N*-isopropylacrylamide) crosslinked by dynamic covalent disulfide bonds. The particles’ swelling is triggered by reduction-induced cleavage of the disulfide bonds into two pendant thiol groups, enabling printing of multilayer scaffolds with high shape fidelity. Without chemically induced swelling, the same formulation requires higher particle concentrations to be printable. Importantly, the printed structures can be annealed by oxidizing the thiol units back into disulfide inter-particle bonds. The printed scaffolds are compatible with human dermal fibroblasts (HDF) and support cell–material interactions. This system offers a unique opportunity for on-demand modulation of both printability and scaffold microstructure, enabling unique control over local stiffness, packing density, and porosity in granular scaffolds.

**Graphical Abstract:** 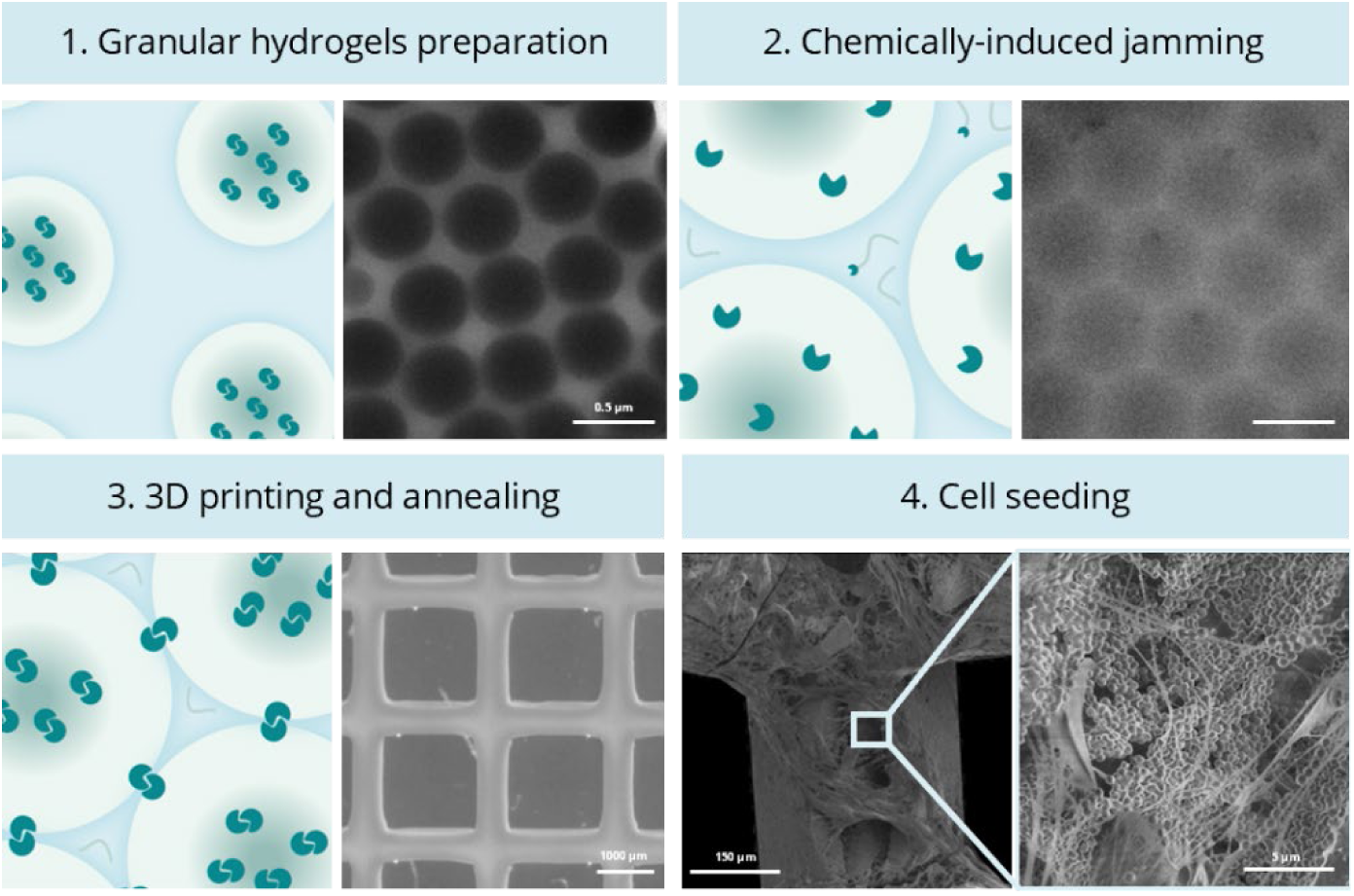

A stimuli-responsive granular hydrogel enables on-demand tuning of yield stress and printability through microgel swelling after jamming. Reductive cleavage of disulfide bonds increases microgel swelling and softness, enhancing packing density and enabling high-fidelity 3D printing. Oxidative annealing restores interparticle bonds, yielding cell-compatible scaffolds with tunable microstructure for tissue engineering.

## 1. Introduction

Granular hydrogels consist of densely packed, so-called “jammed”, hydrogel microparticles that are injectable and exhibit shear-thinning, self-healing, and yield stress behaviors, making them excellent candidates for extrusion-based (bio)printing.^1–3^ Additionally, their inherent porosity, caused by inter-particle voids, facilitates faster cell penetration, migration, diffusion of nutrients, oxygen, and waste, and promotes quicker vascularization of the scaffolds compared to bulk hydrogels.^4, 5^ Furthermore, the ability to alter microgels’ shape, size, and composition enables the creation of diverse granular-based printable materials (biomaterial inks or bioinks) with tunable porosity, mechanical, and physicochemical properties, allowing the rational design of tissue-specific microenvironments that better recapitulate native tissue architecture and function.^6^

The key parameter governing the mechanical and rheological properties of granular hydrogels, as well as their shape retention, is the extent of confinement of individual microgels within a so-called “cage” created by its direct neighbors.^7, 8^ Under these conditions, the collective motion of microgels, which gives rise to macroscopic flow, is restricted by frictional and chemical interactions between densely jammed microgels. Only when applied stress exceeds the yield stress can individual microgels escape their cages, initiating macroscopic flow.^9^ For example, high microgel compaction promotes greater interaction among neighboring microgel polymer chains, increasing the yield stress and thereby facilitating the printing of complex, self-supporting structures without the need for a supporting bath.^10, 11^ Oppositely, low microgel compaction leads to low yield stress and the formation of drop-like filaments with an excess continuous phase, which are unable to withstand the weight of multiple printed layers. Therefore, controlling the yield stress of granular hydrogels is essential for the fabrication of complex three-dimensional architectures. Usually, granular hydrogels with variable yield stress are obtained by dispersing pre-synthesized microgels at varying concentrations (i.e., volume fractions).^12–14^ Under these conditions, the volume of the individual microgels is assumed to remain constant and is dictated by the crosslinking density, if we neglect any deformation caused by high packing density.

Here, we postulate that the microgels do not have to be merely inert building blocks within granular hydrogels, but can instead function as dynamically adjustable volume reservoirs. In such scenario, by modulating the degree of swelling after preparing the granular hydrogel, the yield stress can be adjusted, thereby enabling fine control over material printability and the shape fidelity of the resulting scaffold. As the volume and swelling properties of individual microgels are directly governed by their crosslinking density, crosslinker cleavage represents an attractive strategy for engineering microgels with responsive volume changes. *In situ* swelling can be triggered through the cleavage of dynamic bonds in response to external stimuli, including changes in pH, temperature, light, or redox conditions.^15–20^ Among these dynamic chemistries, disulfide bonds are particularly attractive because they belong to the family of reversible covalent bonds and can be readily cleaved under reducing conditions to generate two thiol moieties.^15, 21, 22^ Interestingly, these thiol moieties can subsequently be oxidized back to reform the original disulfide bonds, enabling reversible modulation of the microgel network. Gaulding et al. elegantly showed the potential of this two-step strategy by transforming jammed yet disconnected poly(*N*-isopropylacrylamide), pNiPAM, microgels internally crosslinked with *N*,*N’*-bis(acryloyl)cystamine (BAC), into cohesive granular hydrogels stabilized by interparticle disulfide bonds.^23^ Building on this concept, we envision an additional advantage for extrusion-based 3D printing. Specifically, the disulfide chemistry can serve a dual function: first, to crosslink the microgels internally, and second, to form interparticle disulfide bonds after printing, thereby annealing and stabilizing the printed construct and preserving its dimensional integrity in aqueous media.

In this study, we present a granular hydrogel system composed of chemically jammed, disulfide crosslinked pNiPAM microgels whose printability can be tuned not only by concentration of particles, but also by their degree of swelling induced through the selective cleavage of the disulfide crosslinks **(Figure 1)**. Using dynamic light scattering, rheology, cryo-electron microscopy, and wet atomic force microscopy (wet-AFM) measurements, we demonstrate that the addition of a reducing agent (tris-(2-carboxyethyl)phosphine hydrochloride, TCEP) effectively cleaves the disulfide bonds, resulting in microgel swelling and partial release of polymer chains. At a fixed weight fraction of particles, this swelling efficiently increases the yield stress of the granular hydrogels, transforming poorly printable formulations into inks allowing to obtain stable, multilayer three-dimensional scaffolds with high shape fidelity. Furthermore, the thiol groups generated upon chemical reduction can be oxidized in a solution of sodium (meta)periodate (NaIO_4_), to form interparticle disulfide crosslinks that impart long-term stability of printed scaffolds immersed in aqueous medium. Finally, the fabricated scaffolds exhibit good cytocompatibility, demonstrating that this strategy provides a versatile platform for the on-demand modulation of printability and the microstructural properties of the resulting granular hydrogel-based 3D constructs.

**Figure 1.**
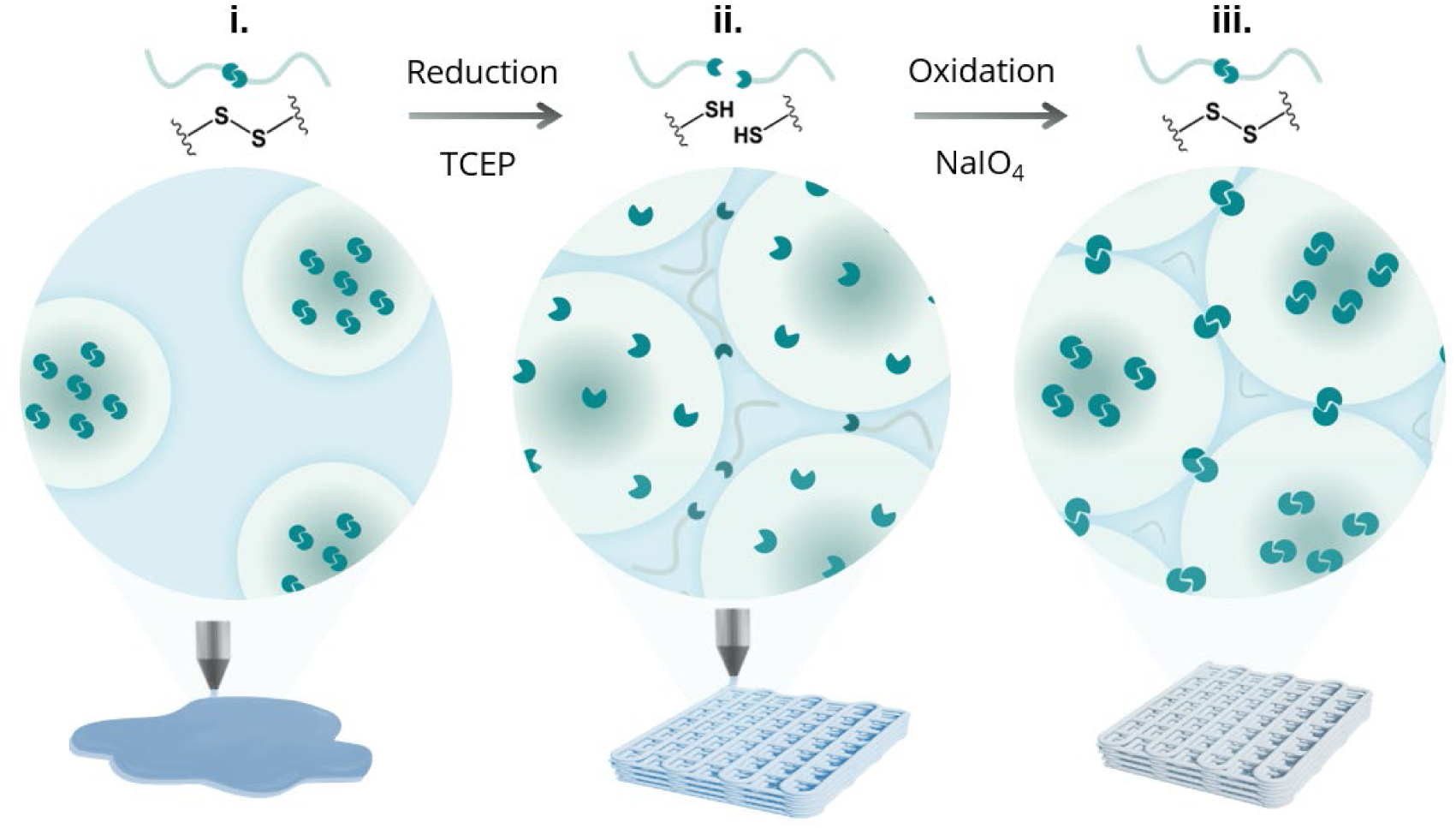
Schematic representation of magnified views of the internal structure of the inks (i and ii) and the printed annealed scaffolds (iii). (i) Pristine granular hydrogels are first prepared at fixed weight fractions and subsequently 3D printed. (ii) The degree of jamming can be increased by inducing microgel swelling through TCEP-mediated reduction of the disulfide crosslinks, enabling the ink to be 3D printed into a stable scaffold. (iii) The printed scaffold is subsequently annealed by immersion in an oxidant solution (NaIO_4_), which promotes the formation of interparticle disulfide bonds and stabilizes the construct.

## 2. Results and Discussion

### 2.1. Microgels synthesis, characterization, and responsiveness to chemical reduction

The pNiPAM based microgels were synthesized by surfactant-free radical dispersion polymerization using a modified procedure from Gaulding et al. (**Figure 2a**).^23^ *N*,*N’*-bis(acryloyl)cystamine (BAC) was used as a crosslinker to straightforwardly introduce the cleavable dynamic covalent disulfide bonds within the network of the microgels during the particle synthesis. The detailed synthesis procedure can be found in the *Experimental section*.

**Figure 2.**
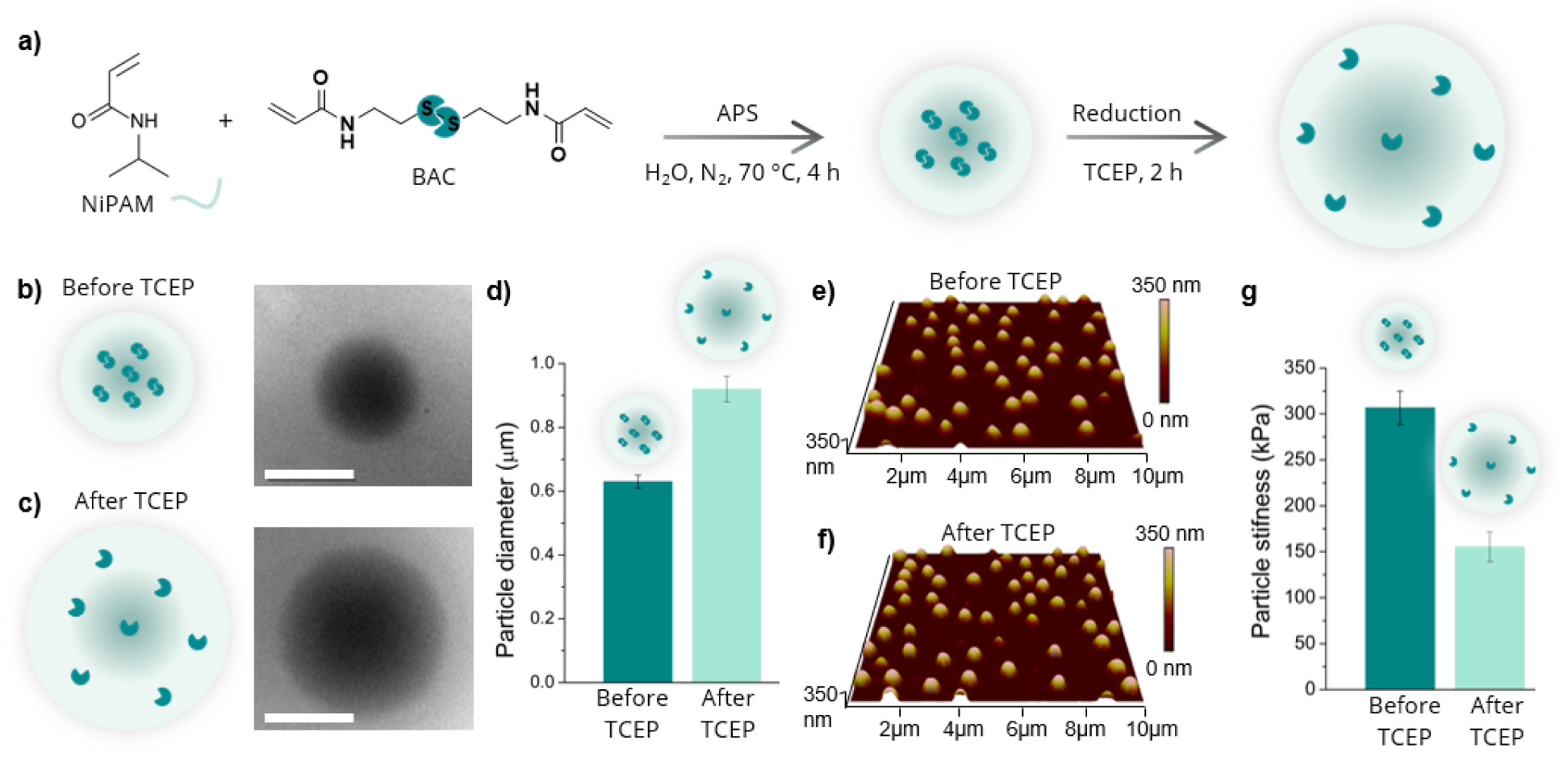
Microgel synthesis and response to TCEP. **(a)** Synthesis scheme of the disulfide crosslinked pNiPAM microgels and swelling induced by chemical reduction with TCEP. Cryo-TEM images of dilute microgel dispersions **(b)** before and **(c)** after addition of TCEP. Scale bars are 0.5 μm. **(d)** Microgels’ average diameter determined from the cryo-TEM image analysis. Wet-AFM images of the height of dilute microgel dispersions **(e)** before and **(f)** after addition of TCEP. **(g)** The microgels’ average stiffness determined by wet-AFM.

Dynamic Light Scattering (DLS) showed that the prepared microgels have an average hydrodynamic diameter of D_H,20°C_ = 0.82 ± 0.03 μm at 20 °C and D_H,45°C_ = 0.5 ± 0.03 μm at 45 °C, indicating narrow size distribution and the typical temperature-induced deswelling behavior of pNiPAM, with Volume Phase Transition Temperature (VPTT) determined to be approximately 32 °C (**Figure S1**). Successful incorporation of BAC, and thus disulfide crosslinks, into the microgel network was confirmed by a combination of ^1^H NMR (**Figures S2-S5)** and UV-Vis spectroscopy titration (**Figures S6).** The BAC content was determined to be 8.3 mol% compared to NiPAM repeat units (*see Experimental section*).

To evaluate the effect of TCEP addition on the swelling behavior and stiffness of individual microgels we performed cryo-TEM, dynamic light scattering (DLS), and wet-AFM measurements under dilute conditions. Cryo-TEM images (**Figures 2b-c** and **Figure S7**) clearly reveal the swelling of the individual microgels following TCEP treatment. The average microgel diameter increased from 0.63 ± 0.02 to 0.92 ± 0.04 μm within 2 hours after TCEP addition (**Figure 2d**). In parallel, DLS measurements highlighted an immediate decrease in the scattering intensity of the microgel dispersion after TCEP addition (**Figure S8**). This observation indicates a decrease in the refractive index difference between microgel and the surrounding medium. We hypothesize that this effect can be attributed to both swelling of the microgels and to the release of loose polymer chains into the surrounding medium.^16, 24^ This release of polymer chains was confirmed after extensive purification by successive centrifugation-redispersion cycles (*see Experimental section*). Briefly, after each centrifugation cycle, the soluble polymer chains were discarded from the sedimented microgel pellet. The cumulative weight loss after purification was 15.0 wt.%, reflecting polymer release induced by TCEP reduction. The effect of TCEP treatment on microgel stiffness was also confirmed by wet-AFM using the PeakForce Tapping Quantitative Nanomechanics (QNM) method (**Figure 2e-g)**.^25^ As expected, pristine microgels (before reduction) showed the highest stiffness (306.7 ± 18.3 kPa). After reduction with TCEP, the stiffness dropped to 155.4 ± 16.1 kPa, which we attribute to the reduced crosslinking density resulting from the cleavage of the disulfide bonds into thiol groups, together with the release of free polymer chains into the interstitial aqueous phase surrounding the microgels. Importantly, the particles did not undergo complete disintegration into free polymer chains, even in the presence of an excess of reductant, as could have been expected based on the fact that the only crosslinker agent used in this study (BAC) is composed of disulfide bonds. This effect can be explained by the commonly reported “self-crosslinking” process resulting from the radical transfer reactions happening during the polymerization of NiPAM and disulfide units, at elevated temperatures.^23^ Therefore, unavoidable and hardly quantifiable inert carbon-carbon and thioether crosslinks are likely formed during the synthesis of the microgels. Unlike disulfide bonds, these crosslinks are not susceptible to reductive cleavage and therefore remain intact after TCEP treatment.

### 2.2. Effect of chemical reduction on the degree of jamming and rheological properties of granular hydrogels

Pristine granular hydrogels were prepared by directly dispersing the microgels in 1x phosphate-buffered saline (1x PBS) at particle weight fractions of 14.0, 16.0, 18.0, 20.0, and 27.8 wt.% (**Table S1)**. More details concerning the preparation procedure can be found in the *Experimental section*. A microgel weight fraction of 27.8 wt.% represented the highest concentration that could be practically achieved by direct dispersion while ensuring homogeneous hydration of the microgel powder. Subsequently, TCEP was added to the granular hydrogels at a 1:1 molar ratio relative to the disulfide crosslinks, thereby inducing microgel swelling, and, consequently, increasing the jamming degree. The rheological properties of the granular hydrogels, before and after the addition of TCEP, were investigated through oscillatory linear frequency sweep and non-linear shear stress sweep measurements (**Figure 3a-b** and **Figure 3c-d**). Interestingly, within the limit of small deformations, i.e., in the linear viscoelastic regime, all pristine granular hydrogels investigated (from 14.0 wt.% to 27.8 wt.%) exhibited a characteristic glass-like behavior, with storage (G’) and loss (G’’) moduli that were nearly frequency independent, where G’ >> G’’ (**Figure 3a**). As the particle weight fraction increased from 14.0 wt.% to 27.8 wt.%, G’ gradually increased from less than 100 Pa to approximately 10000 Pa, reflecting the increasing microgel volume fraction and the corresponding higher degree of jamming. This rheological response is characteristic of a wide range of jammed granular hydrogel systems, irrespective of their chemical composition, in which the high microgel packing density suppresses spontaneous macroscopic flow.^26, 27^ Notably, even for the lowest concentration investigated (14.0 wt.%), the confinement imposed by neighboring particles (“caging”) hindered spontaneous collective motion in the absence of external stress^7, 8^.

**Figure 3.**
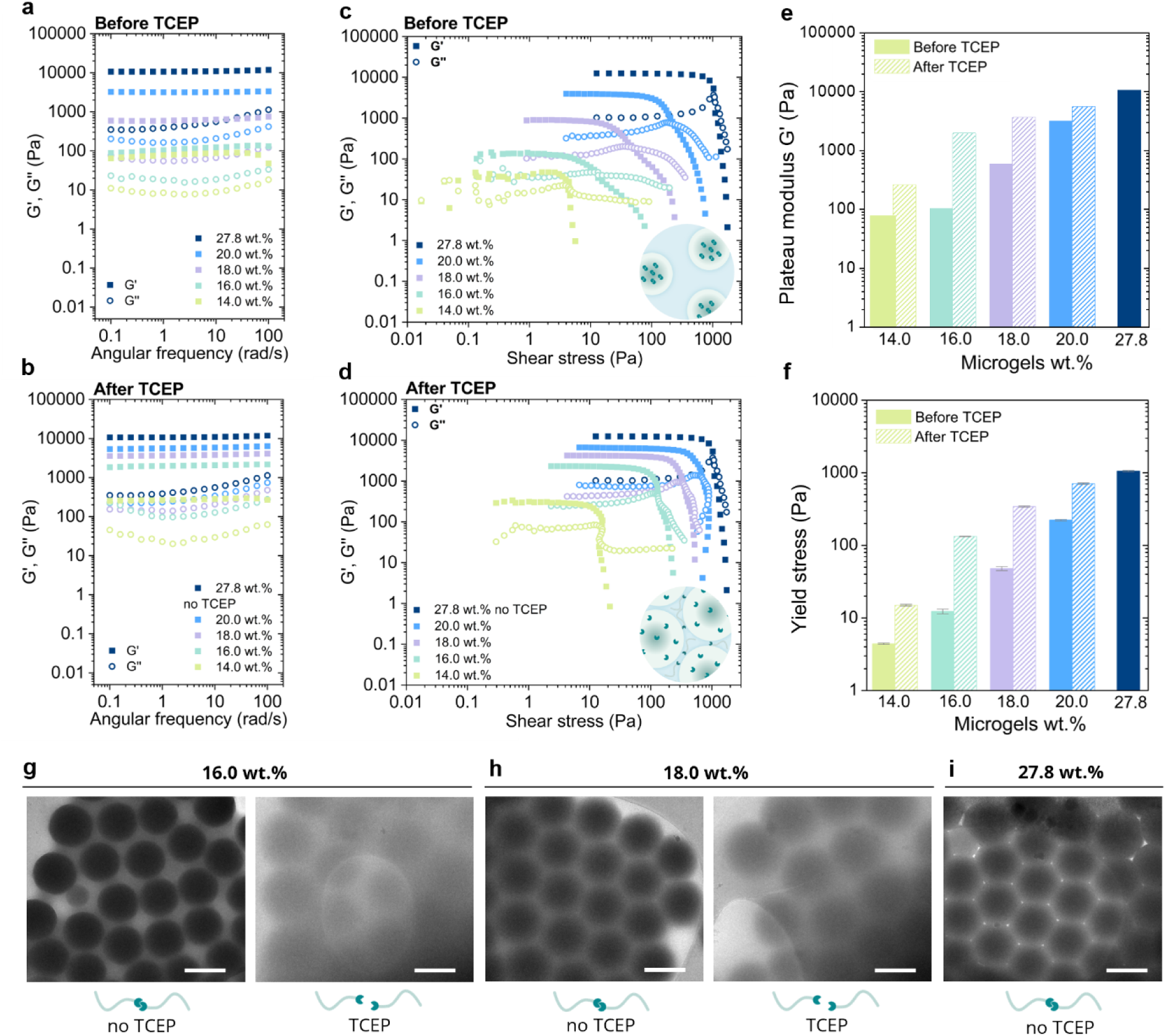
Rheological and structural characterization of TCEP-induced changes in granular hydrogels. Frequency sweep (ω = 0.1 - 100 rad/s, γ = 1% ) **(a)** before and **(b)** two hours after the addition of TCEP. Shear stress sweep **(c)** before and **(d)** two hours after the addition of TCEP. **(e)** Plateau modulus of G’ and **(f)** yield stress of the granular hydrogels before and two hours after TCEP addition for the different microgel concentrations. Yield stress values were determined from the crossover between G’ and G’’ in the shear stress sweeps. Cryo-TEM images of the **(g)** 16.0 wt.%, **(h)** 18.0 wt.%, and **(i)** 27.8 wt.% granular hydrogels at the pristine state (no TCEP added) (left) and two hours after addition of TCEP (right). Scale bars are 0.5 μm.

The ability of these granular hydrogels to flow was further investigated under a gradual increase in shear stress. In **Figure 3c**, the evolution of G’ and G’’ is reported as a function of the applied shear stress experienced by pristine granular hydrogels with concentrations ranging from 14.0 wt.% to 27.8 wt.%. In agreement with the linear viscoelastic data, we observed that at low shear stress values, the granular hydrogels behaved as a non-flowing solids (G’ > G’’) with storage modulus (G’) plateau values gradually increasing from about 30 Pa to 4000 Pa and further to 10000 Pa for the 14.0 wt.%, 20.0 wt.%, and 27.8 wt.% pristine granular hydrogels, respectively. Additionally, above a certain shear stress value, all granular hydrogels exhibited an abrupt drop in G’ concomitant with a kink in G’’. This is the signature of flow initiation in colloidal glass systems, where the “cage”, within which the particles are locked by the neighboring particles, breaks.^7^ From these curves, we extracted the yield stress of the granular hydrogels (**Figure 3f**), i.e., the critical stress value above which macroscopic flow of the material is observed, as hallmarked by the cross-over between G’ and G’’.^26^ As expected, the yield stress also increased gradually with the microgel content from 5 Pa at 14.0 wt.% microgels to slightly above 220 Pa at 20.0 wt.% microgels and to 1050 Pa at 27.8 wt.% microgels. This increase in yield stress reflects the increasing particle volume fraction and, hence, the greater degree of jamming.

The reduction-induced jamming of the microgels was then investigated using the same approach. Directly after the addition of TCEP to the 14.0 wt.% to 20.0 wt.% granular hydrogel inks, the jamming process was followed over time, and both G’ and G’’ increased, reaching a steady-state value after two hours (**Figure S9a**). Then, the frequency sweep (**Figure 3b**) and shear stress sweep (**Figure 3d**) measurements were performed. Similar to the pristine granular hydrogels, the samples that underwent chemical reduction exhibited a classical glass-like behavior with significantly higher G’ and G’’ values. In the low shear stress region, the 14.0 wt.% granular hydrogels showed a 10.0-fold increase in G’ compared to the pristine sample (from 30 Pa to 300 Pa), while the granular hydrogel prepared with 20.0 wt.% microgels showed an increase in G’ from around 4000 to around 7000 Pa (1.6-fold increase). In parallel, all granular hydrogels showed a significant increase in yield stress, reaching 15 Pa (3.0-fold increase) for the 14.0 wt.% sample and 710 Pa (3.2-fold increase) for the 20.0 wt.% sample. Interestingly, the increase upon reduction in the plateau modulus (19.0-fold and 6.2-fold increase) as well as in the yield stress (10.8-fold and 7-fold increase) was most pronounced for the granular hydrogels with particle concentrations of 16.0 wt.% and 18.0 wt.%, respectively. By plotting the evolution of the plateau modulus as a function of the microgels concentration (**Figure S9b**) it is possible to identify two regimes of concentrations where only the hydrogels prepared at 16.0 wt.% and 18.0 wt.% transition after chemical reduction from a loose granular network to a densely packed polymer network that rheologically resembles a homogeneous macrogel network.^28^ This may explain why the hydrogels prepared at these concentrations of microgels exhibit the highest relative increase in elasticity. Overall, the use of TCEP to reduce the microgels disulfide crosslinks to thiol units is an efficient way to increase their jamming extent and consequently increase the yield stress of the resulting granular hydrogels.

To gain further insights into the mechanism by which TCEP induces this greater jamming extent, we employed cryo-TEM to observe the granular hydrogels in native conditions, i.e., at actual jammed microgel concentrations and in the wet state. **Figure 3g-i** and **Figure S10** present cryo-TEM images of the 16.0 wt.%, 18.0 wt.%, and 27.8 wt.% granular hydrogels acquired before and two hours after the addition of TCEP. Before the addition of TCEP, the microgels appeared as spherical beads exhibiting well-defined borders with the surrounding vitreous water, with apparent average diameters of 0.52 ± 0.02 μm, 0.54 ± 0.05 μm, and 0.59 ± 0.04 μm for 16.0, 18.0, and 27.8 wt.% granular hydrogels, respectively. Furthermore, for the 18.0 wt.% and 27.8 wt.% pristine granular hydrogels, the formation of 2D hexagonal packing patterns was observed. This observation suggests that upon increasing the particle concentration, microgels developed slightly faceted edges, a characteristic commonly reported for jammed soft particles^29^. After the addition of TCEP, the grey-level contrast observed in the cryo-TEM images between microgels and surrounding vitreous water drastically decreased. The microgels exhibited a fuzzier structure with clearly less defined borders, with an average size measured in these packed states that significantly increased by ca. 22% (from 0.52 ± 0.02 μm to 0.67 ± 0.03 μm, and from 0.54 ± 0.05 μm to 0.70 ± 0.08 μm for 16.0 wt.% and 18.0 wt.% granular hydrogels, respectively). Overall, the reduction of disulfide crosslinks by adding TCEP led to efficient microgel swelling and softening within granular hydrogels. It resulted in the formation of granular hydrogels in which the degree of microgels jamming was successfully increased, ultimately increasing their yield stress, as revealed by rheological observations.

### 2.3. 3D printing of the granular hydrogel inks

All the proposed granular hydrogel inks, before and after the addition of TCEP, exhibited yield-stress behavior, making them promising candidates for 3D printing. We consequently investigated the effect of particle concentration and the extent of jamming induced by TCEP-mediated cleavage of disulfide bonds on the printability of the resulting granular hydrogels.

First, the extrudability, i.e., the ability to form continuous filaments, of granular hydrogels prepared at a fixed particle weight concentration, before and after reduction-induced jamming, was assessed (**Figure 4**).^3^ Before TCEP addition, inks with lower microgel concentrations (14.0 wt.% and 16.0 wt.%) were extruded as droplets or short droplet-like fibers (“pear-shaped”). In contrast, inks prepared with higher microgel concentrations (18.0 wt.% – 27.8 wt.%) were extruded as continuous, well-defined filaments. Strikingly, after TCEP addition, the 16.0 wt.% – 27.8 wt.% inks were extruded as continuous filaments, irrespective of microgel concentration. Generally, higher microgel concentration inks required higher extrusion pressures to achieve uniform fiber formation. Additionally, the chemically-jammed inks obtained after the addition of TCEP also required higher extrusion pressures compared to inks with the corresponding pristine granular hydrogel inks (for example, the extrusion pressure increased from 40 kPa to 70 kPa for the 20.0 wt.% ink). This observation is explained by their increased yield stress, resulting from increased particle volume fraction and consequently more interparticle contact.

**Figure 4.**
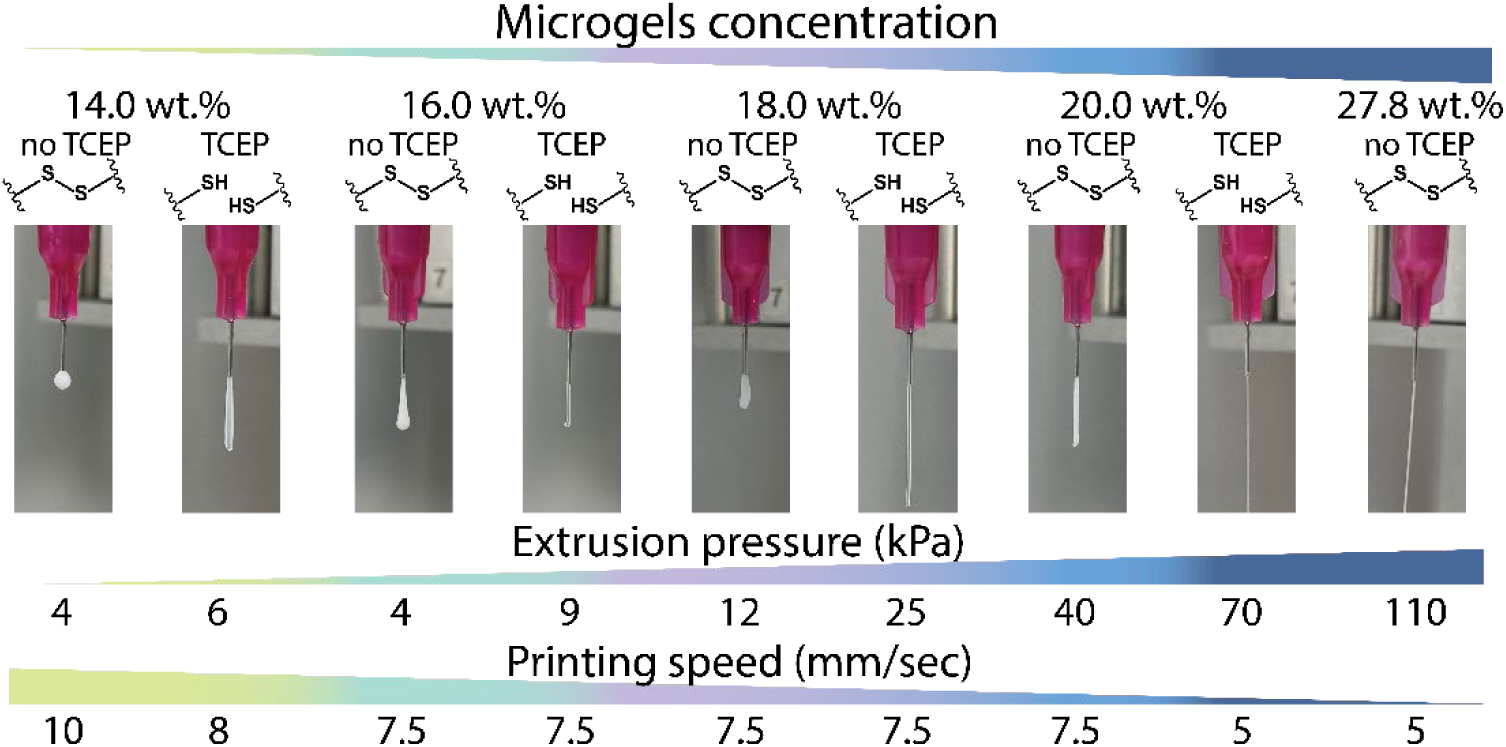
Extrudability of the different granular hydrogel inks. Granular hydrogel inks extrusion at increasing microgel concentration (14.0 wt.%, 16.0 wt.%, 18.0 wt.%, 20.0 wt.%, and 27.8 wt.%) before (pristine) and after reduction-induced jamming with TCEP. Extrusion pressure in kPa used to extrude a filament of each ink and printing speed used for square-mesh printing in mm/sec.

To further evaluate the stacking ability and shape fidelity of the granular hydrogel inks, multilayered square-mesh scaffolds (2, 4, 18, 16, and 32 layers) were 3D printed by pneumatic-driven extrusion (**Figure 5**). The corresponding printability indexes (Pr), reported in **Figure S11**, were calculated using Equation 1 described in the *Experimental Section*. The Pr index quantifies how close the pore shape is to the ideal square geometry of the designed pattern^30^. A Pr value of 1 reflects square pores and, consequently, high print fidelity. Pr values higher than 1 represent irregular extrusion (over-gelation). Conversely, Pr values smaller than 1 indicate fusion or spreading of the printed filaments (under-gelation). Before TCEP addition, the 14.0 wt.% ink exhibited poor shape fidelity even when only 2 layers were printed, with spontaneous flow observed at the intersection between strands, leading to a Pr value of 0.86 ± 0.01. As the number of layers increased, adjacent strand merging due to ink flow led to the lowest Pr value, regardless of the number of printed layers. As the microgel concentration increased to 16.0 wt.% and 18.0 wt.%, the shape fidelity, and consequently the Pr value, increased regardless of the number of printed layers, compared to the 14.0 wt.% ink. Nevertheless, diminished shape retention and pore geometry loss were observed when printing 8 layers (Pr = 0.80 ± 0.01 and Pr = 0.82 ± 0.01 for microgels concentrations of 16.0 wt.% and 18.0 wt.%, respectively) or more. For the inks prepared at higher microgel concentrations (20.0 wt.% and 27.8 wt.%), increasing the number of printed layers did not cause deformation or shape relaxation of the bottom layer. Instead, the scaffold pores remained nearly square even after printing 16 layers (Pr = 0.87± 0.01 and Pr = 0.90 ± 0.01, respectively), demonstrating the superior shape retention of these inks, consistent with the highest yield stress values.

**Figure 5.**
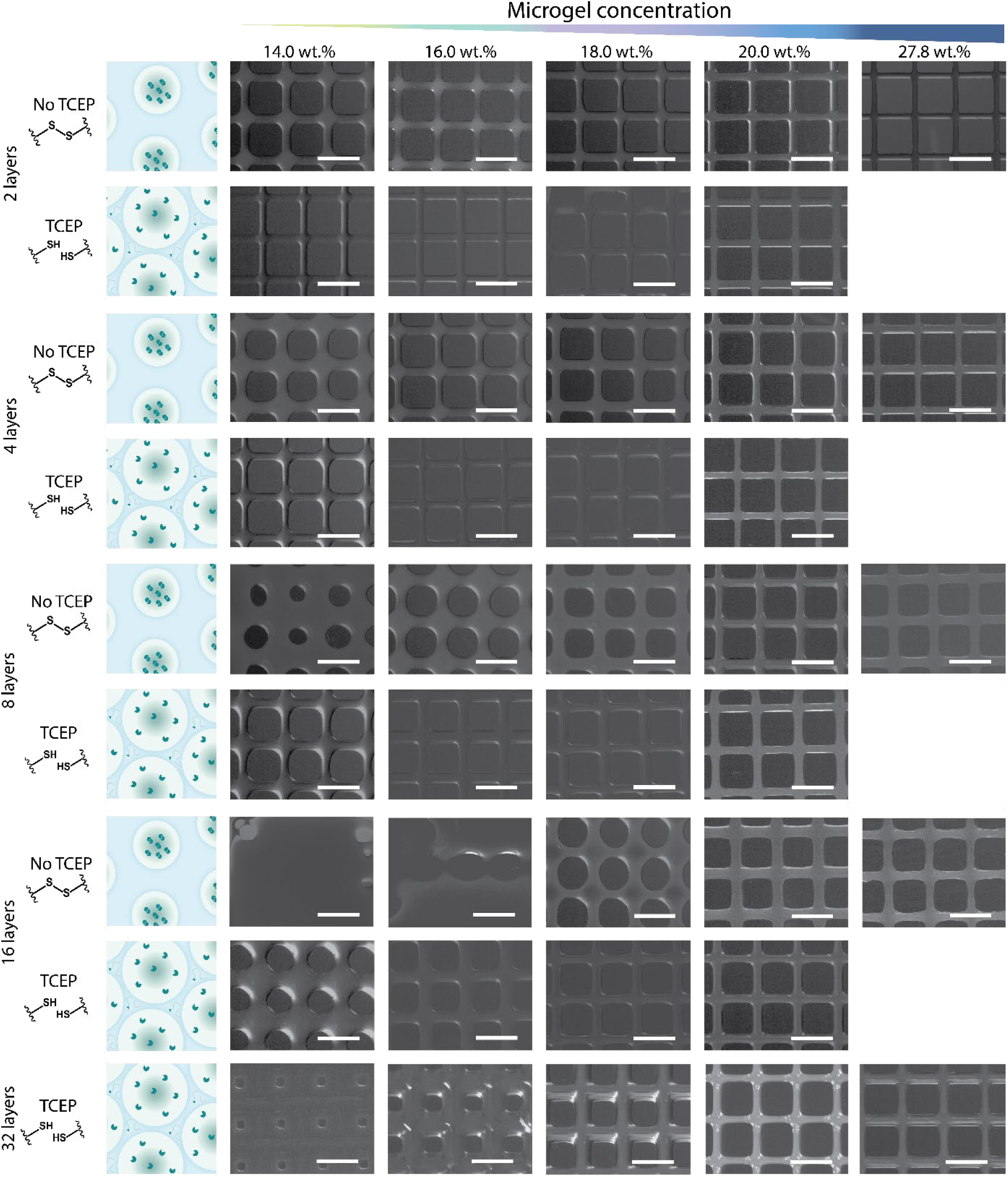
3D printing of the granular hydrogel inks. Optical images of the 3D printed square mesh scaffolds obtained by pneumatic-driven extrusion of the granular hydrogel ink at varying concentrations. Each concentration from 14.0 wt.% to 20.0 wt.% was printed before (pristine state) and after TCEP addition into 2-, 4-, 8-, 16- and 32-layered constructs. Scale bars are 2000 µm.

The addition of TCEP and subsequent microgel swelling and reduction-induced jamming improved printability of the granular hydrogel inks and shape retention of printed scaffolds, as evidenced by the Pr increase, irrespective of microgel concentration, compared to granular hydrogel inks before TCEP. Similarly to what was noticeable with the granular hydrogel inks before reduction, increasing microgel concentration led to better shape fidelity and printability. For the ink with the lowest microgel concentration, i.e. 14.0 wt.%, the effect of disulfide reduction by TCEP and subsequent reduction-induced jamming was already visible when printing 2-layered scaffolds. In this case, the microgels’ swelling enabled doubling the number of printed layers compared to the pristine ink. As the microgel concentration increased to 16.0 wt.% and 18.0 wt.%, the effect of reduction-induced particle jamming was most prominent, which aligns with the highest relative increase of the rheological characteristics reported earlier. Pristine granular inks with 16.0 wt.% showed low shape retention even when printing 2 layers. However, after TCEP addition, the same ink was printed into dimensionally stable 8-layered scaffolds, four times taller than what was possible before chemical reduction. Similarly, the 18.0 wt.% granular ink exhibited a higher yield stress and could therefore withstand 4 printed layers with superior Pr values. However, 8-layered prints were not successful due to material relaxation and flow. After TCEP addition, the same ink enabled the fabrication of 16-layered scaffolds with high Pr values, again representing a four-fold increase in printable height compared to the pristine granular hydrogel. The 20.0 wt.% ink, before and after TCEP, showed the highest Pr value compared to the inks with lower microgels concentrations, regardless of the number of printed layers. Also, no significant change in the Pr value was observed for the 20.0 wt.% granular hydrogels after TCEP addition, when printing thicker scaffolds up to 16 layers. Therefore, at this microgel concentration, reduction-induced jamming was not critical for obtaining dimensionally stable scaffolds up to 16 layers thick. Interestingly, the largest improvement in printability was observed for the 16.0 wt.% and 18.0 wt.% granular hydrogels, in agreement with the rheological and cryo-TEM analyses, suggesting that the transition from a loosely to a densely packed granular network governs the printability of these granular hydrogel inks.

Overall, the concentration and the extent of microgel jamming directly dictate the yield stress value and, as a consequence, shape fidelity of the printed scaffolds. When the microgel concentrations were below 20.0 wt.%, the effect of the reduction-induced jamming by TCEP on the printability was significantly high, consistent with the corresponding increase in the yield stress. Notably, the effect of chemical reduction became increasingly pronounced when printing thicker scaffolds. Below 20.0 wt.% microgel concentration, swelling of the individual particles was efficient in creating a space-filling network that increased yield stress into a range enabling the fabrication of dimensionally stable scaffolds up to 16 layers high. In contrast, for the pristine ink prepared with 20.0 wt.% microgels, the yield stress was already sufficiently high to obtain stable scaffolds up to 16 layers. Consequently, although TCEP-induced microgel swelling further increased the yield stress, this did not translate into a noticeable improvement in the shape fidelity of the printed scaffolds.

### 2.4. Annealing of the 3D printed scaffolds and stability in physiological conditions

For practical use in tissue engineering, the scaffolds must remain dimensionally stable, i.e., neither dissolve nor deform under physiological conditions, at least during the time frame required for the cell to adhere and proliferate.^31^ For granular hydrogels, a usual strategy to achieve scaffold stability is to perform an annealing reaction, which consists of creating stable chemical or physical bonds between neighboring particles.^26^ Here, we wanted to take an advantage of the reversibility of dynamic chemistries, specifically by forming disulfide interparticle bonds from the newly generated thiol moieties by oxidation with a strong oxidizing agent such as NaIO_4_ (**Figure S12**)^23^. Therefore, to proof this possibility, we immersed printed scaffolds with or without annealing ( i.e. incubating in a 10^−2^ M NaIO_4_ solution for two hours) in 1x PBS. The average strand thickness as a function of immersion time at 21 °C at both conditions is shown in **Figure 6a**. Without annealing, the scaffold spontaneously disassembled, and the microgels dispersed in the medium. After annealing, however, the scaffolds remained dimensionally stable for at least 14 days (longer time periods were not tested), with an average strand thickness of 507 ± 96 μm (compared to an initial strand thickness of 476 ± 45 μm) **(Figure 6a)**. The stability of the printed scaffold microstructure was also examined using cryo-SEM (**Figure 6b** and **Figure S13**). Before annealing (in air), the microgels appeared closely packed and showed a relatively distorted circular shape, highlighting their deformability after TCEP-induced chemical reduction (**Figure 6b/iii**). Notably, after annealing, the microgels contracted significantly and regained a more circular shape, resulting in the formation of voids and limited contact points between neighboring particles (**Figure 6b/iv**).

**Figure 6.**
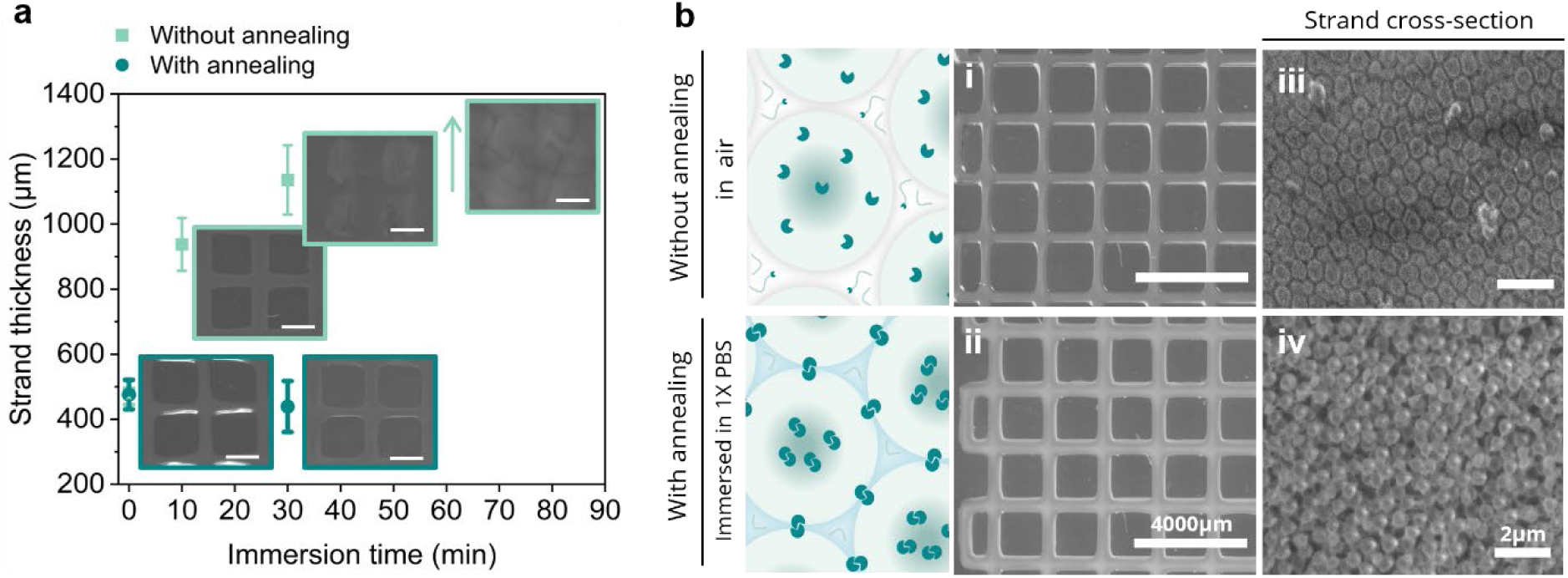
Annealing of the 3D printed scaffolds. **(a)** Strand thickness as a function of the immersion time in 1x PBS with and without annealing. Scale bars are 2000 µm. **(b)** Optical microscopy images of a 4-layered 3D printed scaffold (20.0 wt.% after TCEP addition) (i) in air without annealing, and (iii) after annealing with NaIO_4_ and immersion in 1x PBS at 21°C. Cryo-SEM images of the cross-section of a 3D printed strand of the 20.0 wt.% after TCEP addition (iii) in air without and (iv) with annealing with NaIO_4_ and after subsequent immersion in a 1x PBS solution.

The contraction of the microgels upon oxidation with NaIO_4_ was further confirmed by DLS, which showed a decrease in the average hydrodynamic diameter of the microgels from 897 ± 12 to 868 ± 32 nm at 20 °C (**Figure S14a**), and by wet AFM, where the microgel stiffness increased from 155.4 ± 16 to 246 ± 21 kPa (**Figure S14b-c**). Overall, we hypothesize that the annealing-induced particle contraction is caused by the re-formation of a significant number of intraparticle disulfide crosslinks, and combined with the formation of interparticle disulfide crosslinks that permanently connect neighboring microgels and stabilize the scaffold under wet conditions.

The proposed strategy is promising for controlling the long-term dimensional stability of scaffolds using the same crosslinking chemistry both within and between the microgels. It also induces internal porosity formation within the scaffold, which may further enhance their cytocompatibility by facilitating a more efficient exchange of nutrients, oxygen, and metabolic waste with the surrounding culture medium.^31, 32^

### 2.5. Cell culture studies

To evaluate the biological performance, the 20.0 wt.% formulation was chosen, as it showed the best printing performance, shape retention, and long-term stability, therefore being the most suitable candidate for evaluating the cytocompatibility of the proposed system **(Figure 7)**. To start, the indirect cytocompatibility of the chemically reduced granular hydrogel inks was assessed. To this end, primary human dermal fibroblasts (HDFs) were seeded in tissue culture plates, and a printed strand of annealed granular hydrogel was subsequently immersed in the culture medium. Cell viability was evaluated using live/dead staining after 3, 7, and 14 days of culture. As shown in **Figures S15a** and **S15d,** no significant cytotoxicity was observed throughout the study, with cell viability consistently exceeding 90%, indicating that the annealed granular hydrogel does not adversely affect cell survival over 14 days of culture.

**Figure 7.**
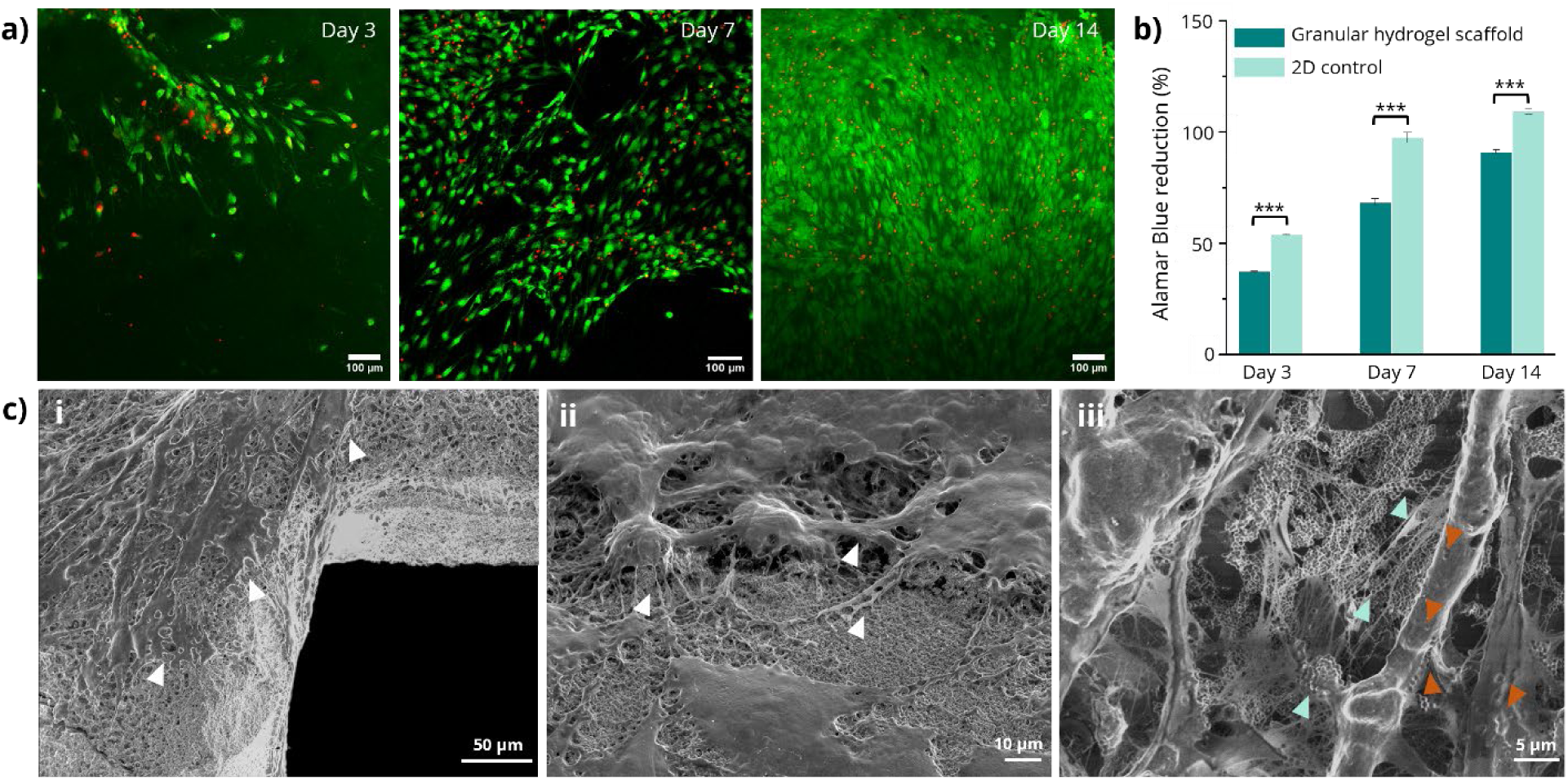
HDF cells – granular hydrogel scaffold interactions. **(a)** Live/dead assay and **(b)** metabolic activity of HDF cells seeded on top of the printed granular hydrogel scaffolds (microgel concentration of 20.0 wt. %) and 2D control (cells seeded on 24-well plates) after 14 days of culture. **(c)** Cryo-SEM of the printed scaffolds after 14 days of HDF cells cultured on top. Cell spreading (white arrows), cell pulling and surface reorganization (blue arrows), possible microgel internalization (red arrows).

Subsequently, the cytocompatibility under direct cell-material contact conditions was investigated by seeding HDFs onto the annealed granular hydrogels. For these experiments, the hydrogel ink was first deposited to fully cover the bottom surface of the culture wells (without printing), then annealed, washed with 1 x PBS, and incubated overnight in fetal bovine serum (FBS) at 37 °C before cell seeding. After 14 days of culture, live/dead staining was performed (**Figure S15b**). Importantly, throughout all experimental procedures, including staining and confocal imaging, the temperature was carefully maintained to preserve the thermoresponsive state of the material and prevent cell detachment. Confocal microscopy revealed the formation of a nearly continuous cellular monolayer across the granular hydrogel surface after two weeks of culture, where the vast majority of the cells were alive (76.9 ± 14.9 % viability) (**Figure S15bi** and **S15d)**. Notably, while the cells exhibited widespread surface coverage, pronounced cell–cell interactions also became evident, resulting in the formation of interconnected cellular clusters that appeared to dominate over cell–material interactions (**Figure S15b/ii-iv**). This can result from cells pulling on the crosslinked microgels, and slowly detaching from the rest of the casted granular hydrogel. In the cases of the cell sheet detachment from the jammed microgels scaffold, more dead cells were observed.

Having confirmed both indirect and direct material cytocompatibility, we evaluated the ability of the printed scaffolds to support long-term cell growth. Therefore, grid-like architectures were 3D printed, followed by annealing, washing, and overnight incubation in FBS at 37 °C before HDF seeding. Cell viability and metabolic activity were assessed on days 3, 7, and 14 using live/dead staining and alamarBlue assay, respectively. The printed granular hydrogels supported both short- and long-term cell viability, with viabilities of 68.5 ± 2.6 % and 91.3 ± 4.8 % on days 3 and 14, respectively **(Figure 7a)**. Furthermore, image analysis revealed a progressive increase in cell density throughout the culture period, indicating sustained cellular proliferation. This observation was corroborated by the continuous increase in metabolic activity from 37.4 ± 0.2 % to 91.1 ± 0.9 %, confirming that the cells remained metabolically active and proliferative over the 14 days **(Figure 7b)**. The HDF cells were well spread, and fewer detaching clusters were observed compared to the cast microgels. We hypothesize that this is due to the higher jamming degree of printed granular hydrogels than the casted ones, caused by the applied extrusion pressure. As a consequence, the interparticle crosslinks were more pronounced, which hindered not only the pulling on single microgel by the cells, but also the infiltration trough the scaffold as a result of a possible change in void fraction.

To deeply investigate cell distribution and interactions within the printed scaffolds, cryo-SEM was performed after 14 days of culture. As a comparison, printed scaffolds immersed in complete media for 14 days but without cells were also imaged (**Figure S15ci**). As shown in **Figure 7c/i-ii**, HDFs exhibited a well-spread morphology characterized by numerous elongated cytoplasmic protrusions extending along and between the jammed microgels (white arrows). Cells formed an interconnected network throughout the scaffold, demonstrating extensive spreading and strong intercellular connectivity, bridging the printed strands of the scaffolds. Cryo-SEM images also showed cell-to-microgel connections, with clear interactions between the fibroblasts and single microgels (**Figure 7c/iii**, blue arrows). The material-compliant mechanical properties and the HDFs natural mechanobiology, characterized by contractile forces, facilitated local matrix rearrangement. Additionally, HDF cells were likely internalizing some microgels, as shown by the red arrows in **Figure7c/iii** and **Figure S15c/vi,** visible as local changes in the surface morphology and smoothness of the cell. Via microgel pulling or/and internalization, the cells were breaking connections with the rest of the bulk scaffold (blue and red arrows, respectively), which most probably led to single microgel detachment (**Figure 7c/iii** and **Figure S15c/ii-vi**).

Overall, these findings demonstrate that dynamic granular hydrogels can simultaneously provide excellent printability, long-term structural stability, and a permissive microenvironment for cell growth. Together, these properties highlight the promise of this class of responsive granular hydrogel inks as versatile platforms for investigating cell–microgel interactions and for developing advanced biomedical systems, including cell-instructive scaffolds, cellular uptake platforms, and controlled therapeutic delivery vehicles. Future studies should further elucidate how microgel size, architecture, and dynamic remodeling influence cell attachment, infiltration, and interactions with the 3D-printed scaffolds.

## 3. Conclusion

In this study, we have demonstrated the unique capability to use a chemical trigger to control, in situ, the extent of microgel jamming, which dictates the yield stress properties of the resulting granular hydrogels and their subsequent 3D printability. We have shown that disulfide-crosslinked pNiPAM microgels can swell and release free polymer chains after reduction with TCEP. The combination of swelling and polymer chain release enabled the use of the microgels as a volume reservoir to fill the interstitial space between them, thereby further increasing both the yield stress and the shape-retention ability of the resulting granular hydrogels. Inks with a microgel concentration of 18.0 wt.% could be printed into square-mesh scaffolds that exhibit very high printability up to 32 layers. In contrast, at the same particle concentration before the reduction-induced jamming, it was only possible to print 4 layers without observing material flow. Post-printing annealing of the microgels was achieved by oxidizing back free thiols into inter- and intra-particle disulfide bonds. This dynamic thiol–disulfide chemistry therefore served a dual function: it enabled reduction-induced microgel swelling and enhanced printability before 3D printing, while subsequent oxidation stabilized the printed scaffolds under physiological conditions. Tuning the balance between particle interactions governs material flow, strand microstructure, and scaffold integrity, making thiol-crosslinked granular inks highly promising for advanced tissue engineering and biofabrication with improved control. The system proved to be cytocompatible, favoring cell attachment, spreading, and proliferation. Altogether, this work demonstrates how dynamic covalent chemistry in granular hydrogel systems can be harnessed to control printability, scaffold stability, and cellular performance simultaneously. Importantly, the chemically induced swelling by disulfide bond cleavage can be applied to other microgel types, in terms of size, chemical composition, and even shape, to develop tunable granular hydrogels with in situ control over the final scaffold microstructure and mechanical properties, leading to increased scaffolds (local) heterogeneity. Therefore, this work opens new opportunities for the development of next-generation versatile and controllable granular inks for biomimetic 3D printing and biofabrication.

## 4. Experimental Section

### 4.1. Materials and reagents

*N*-isopropylacrylamide (NiPAM, > 99%), ammonium persulfate (APS, > 99%), and sodium (meta)periodate (NaIO_4_, > 99%) were purchased from Sigma-Aldrich. *N*,*N’*-bis(acryloyl)cystamine (BAC, 98%), tris-(2-carboxyethyl)phosphine hydrochloride (TCEP, > 99%), 5,5-dithio-bis-(2-nitrobenzoic acid) (DTNB), and 1x phosphate-buffered saline (1x PBS, D8537, Sigma-Aldrich, the Netherlands) buffer solution were purchased from Thermo Fisher Scientific. All chemicals were used as received. Reverse osmosis water with a minimum conductivity of 10 μS.cm^−1^ was used for all experiments.

### 4.2. Microgels synthesis

The microgels were synthesized by surfactant-free precipitation polymerization. 7.05 g of NiPAM (62.3 mmol) and 0.81 g of BAC (3.10 mmol, 5 mol% compared to NiPAM monomer) were dissolved in 690 mL of water in a round-bottom flask under stirring at ambient temperature and under N_2_ flow for 40 minutes. The reaction medium was then heated to 70 °C under N_2_ flow for 30 minutes. Subsequently, 0.34g of APS (1.27 mmol, 2 mol% vs. NiPAM monomer), previously dissolved in 10 mL of water, was injected into the reaction mixture, and the reaction was allowed to progress for four hours under continuous stirring and N_2_ flow. Finally, the reaction medium was exposed to air, and the flask was immersed in an ice-cold bath to stop the polymerization. The microgels were purified from unreacted monomers and free polymer chains by 5 consecutive cycles of centrifugation-redispersion (30 000 rpm for 15 minutes at 4 °C, 50.2 Ti rotor from Beckman Life Sciences) in deionized water. After purification, the microgels were freeze-dried and stored at room temperature until further use.

### 4.3. Determination of the BAC content in the microgels

The BAC content in the microgels was determined by colorimetric titration of the thiol moieties resulting from the reduction of the BAC by TCEP with DTNB^23^. First, the reaction between BAC and TCEP in a 1:1 mol ratio was confirmed to be quantitative (> 95% yield after 15 min) by ^1^H NMR (**Figure S2-S5**) in deuterated 1x PBS (pH 7.2). Then, 2 mg of (dry) microgel powder was dispersed in an 8 mL 1x PBS solution, and 2.2 mg of TCEP was further added as powder to reach a concentration of 10^−3^ M (big excess compared to the supposed BAC content). Next, 3.1 mg of DTNB (10^−3^ M) was added to allow for the reaction with the thiol groups, which yielded the production of the yellowish 2-nitro-5-thiobenzoic acid (TNB) product that strongly absorbs at 408 nm. The molar extinction coefficient of the TNB created by this reaction pathway (BAC + TCEP + DTNB successively) was determined in parallel with these molecules (no microgels involved) to be 16 180 L.mol^−1^.cm^−1^ (**Figure S6a** and **S6b**). Finally, the yellowish microgel dispersion was ultracentrifuged to avoid any interference coming from the turbidity of the sample, the supernatant was retrieved and diluted by 5, and measured by UV-Vis spectroscopy (**Figure S6c**). An absorption peak at 408 nm was observed with an absorption value of 0.934. The BAC content was determined to be 8.3 mol% compared to NiPAM repeat units (see **Table S2** for the calculations).

### 4.4. Dynamic light scattering (DLS)

DLS was performed at a detection angle of 90°, using a Zetasizer (Ultra Malvern Instrument, United States of America) equipped with a HeNe laser (λ = 632.8 nm). Before DLS measurements, all samples were prepared with deionized water or 1x PBS at a microgel concentration of 0.1 mg.mL^−1^ (0.01 wt. %). The microgel solutions were equilibrated overnight at room temperature. To study the variation of the size of the microgel as a function of temperature, the samples were measured from 10 to 45 °C every 1 °C with 10 min temperature equilibration between each measurement. All analyses were performed with the software supplied by the manufacturer. The results were given as intensity-averaged hydrodynamic diameters (mean diameters based on the intensity of the scattered light). Three replicates were performed for each temperature. The reduction-induced swelling of the microgels by addition of TCEP at a concentration of 10^−2^ M was followed over time by measuring the average hydrodynamic diameter and the absolute scattering intensity every 90 seconds after addition of TCEP at a temperature of 20 °C. The scattering intensity data is reported as a relative value compared to the absolute scattering intensity at time 0 before the addition of TCEP.

### 4.5. ^1^H NMR

^1^H NMR spectra were acquired on an Agilent 400-MR 400 MHz spectrometer at 25 °C. Samples were dissolved in deuterated 1x PBS and measured with a pulse width of 45 μs, spectral width of 12/-2 ppm, recycle delay of 1 s, and 32 scans. Chemical shifts were determined from tetramethylsilane referenced to the residual isotopomer solvent signal (HOD).

### 4.6. UV-Visible spectroscopy (UV-Vis)

The optical absorbance measurements were carried out with a UV–Vis Hitachi U-1800 spectrophotometer using a 1 cm path length quartz cell, in a wavelength range from 200 to 800 nm at 20 °C.

### 4.7. Cryo-transmission electron microscopy (cryo-TEM)

The cryo-TEM experiments were performed by placing 2 µl of the microgel dispersions (diluted) and granular hydrogels on a holey carbon supported copper grid (Plano, Wetzlar, Germany, type S147-4) that was further blotted for 2 s and vitrified in undercooled liquid ethane using a Gatan (Pleasanton, OR, United States) CP3 plunge-freezer. The samples were transferred to a Gatan model 914 cryo-TEM sample holder under liquid nitrogen and visualized using a JEOL (Akishima, Tokyo, Japan) JEM-2100 LaB_6_ TEM at 200 kV accelerating voltage under low-dose conditions (CCD camera Gatan Orius SC1000, 2 s acquisition time). The mean diameter was obtained from measuring 30 particles.

### 4.8. Determination of the microgel weight loss after reduction with TCEP

The content of polymer chains released from the crosslinked microgel structure was measured gravimetrically. First, 31.8 mg of microgels were dispersed in 4 mL of 1x PBS. After complete dispersion, 4.5 mg of TCEP (1:1 mol ratio vs BAC) was added, and the mixture was left under stirring conditions for two hours. Then, the dispersion was subjected to 4 cycles of centrifugation-redispersion against deionized water (30 000 rpm for 10 minutes at 4 °C, MLA 80 rotor from Beckman Life Sciences) to remove free polymer chains, residual TCEP, and salt. The complete removal was reached when the conductivity of the supernatant was equal to the conductivity of deionized water (<10 µS.cm^−1^). Finally, the microgel pellet was dried at 50 °C overnight to obtain 27.0 mg of dry polymer content. The relative polymer weight loss upon reduction with TCEP was then determined to be (31.8 - 27.0) / 31.8 = 15.1 wt.%.

### 4.9. PeakForce Tapping Quantitative Nanomechanics (QNM)

First, borosilicate glass substrates (10 mm × 10 mm) were cleaned by sonicating for 15 minutes in soapy water, water, acetone, and isopropanol. Then, a layer of chromium (5 nm) and gold (40 nm) were thermally evaporated on top of the clean substrates. Afterward, monodisperse pNiPAM microgel solutions (1 mg/mL in PBS), pristine, treated with TCEP, and treated with TCEP followed by NaIO_4_ treatment, were deposited (50 µL) onto the gold substrates. The substrates were allowed to dry overnight at room temperature, facilitating non-covalent microgel attachment. The surfaces were then rinsed to remove unbound microgels before the AFM measurements. The softness of individual microgels at the pristine state, after reduction, and after reduction followed by oxidation, was measured using PeakForce Quantitative NanoMechanics (QNM) mode on an AFM (Dimension 3100 Nanoscope V, Bruker, Germany) in the liquid state. The samples were scanned at a peak force of 40 nN using a SCANASYST-FLUID probe (nitride-coated silicon, Bruker, Germany) featuring a v-shape nitride cantilever with a nominal spring constant of 0.7 N/m, measured by the software. The oscillation frequency of the Z-piezo was 0.3 kHz, and the peak force amplitude was set to a default value of 10 μm. The force–displacement curves for the AFM image have a resolution of 128×128 pixels. All measurements were performed at room temperature. Individual particle softness was measured with NanoScope Analysis 1.8 software (Bruker). For that, the average of the Sneddon Modulus was taken from a 329.22 x 329.22 μm square of 30 particles per sample.

### 4.10. Granular hydrogels preparation

The microgels were dispersed in 1x PBS at concentrations of 14.0 wt.%, 16.0 wt.%, 18.0 wt.%,

20.0 wt.% and 27.8 wt.% to form granular hydrogels. To obtain homogeneously dispersed microgels, three consecutive cycles of temperature change were performed under stirring: heating to 28 °C followed by cooling to 20 °C. The increase in temperature allows microgel partial deswelling (staying below the LCST of pNIPAM to avoid microgels aggregation), leading to a lower effective volume fraction and consequently more liquid dispersion, making it easier to practically disperse particles in 1x PBS. Those granular hydrogel inks, so-called pristine, were used for rheology and cryo-TEM characterisation, and 3D printing. To obtain the reduction-induced jamming of the microgels, TCEP was added directly as powder under mechanical stirring at a 1:1 mol ratio compared to BAC, at 20 °C (see **Table S1**). After two hours, the “after TCEP addition” experiments were performed, including rheology, cryo-TEM, and 3D printing. The two-hour waiting time required for complete reduction-induced jamming to be completed was determined via time sweep rheological experiments (see **Rheology** section).

### 4.11. Rheology

An Anton Paar MCR302 rheometer (Anton Paar Co., Netherlands) with a cone-plate geometry (cone angle 1°, cone diameter 25 mm) was used to perform small amplitude oscillatory shear rheology. The analyses were performed at a controlled temperature of 20 °C. Around 0.2 mL of the granular hydrogels was collected using a spatula and loaded on the bottom plate of the rheometer. The gap was fixed at 49 µm, and then a low viscosity oil (Fluka silicon oil, viscosity 50 mPa.s) was added around the geometry to prevent solvent evaporation, unless otherwise stated. After the normal force decreased and stabilized to a constant value (between 0 and 1 N depending on the microgel concentration), a strain sweep (strain γ = 0.1 to 1000 %, angular frequency ω = 1 rad/s) was performed to confirm the materials’ linear viscoelastic regime (LVE) and study the yield stress properties. Then, frequency sweep measurements (from ω = 0.1 to 100 rad/s at γ = 1%) were performed to measure relative elastic (G’ storage modulus) and viscous (G’’ loss modulus) components. The in-situ reduction-induced jamming was followed just after the addition of TCEP by performing a time sweep experiment (ω = 1 rad/s at γ = 1%) for 6 000 s. Frequency and strain sweeps were performed on the TCEP-reduced samples as described above for pristine samples.

### 4.12. 3D Printing

For 3D printing, the granular hydrogels were loaded into a 10 mL cylindrical syringe and centrifuged at 1000 rcf for 3 min to eliminate possible air entrapped. A 25G standard blunt needle (0.25 mm inner diameter) was connected to the cartridge used for printing. Scaffolds were fabricated using a pneumatic-driven extrusion 3D printer (Bioscaffolder 3.3, GeSiM mbH, Radeberg, Germany). Different printing speeds and printing pressures were used to obtain a well-defined printed structure. Strand height and width were kept at 0.050 mm. Scaffolds with 2, 4, 8, 16, and 32 layers were printed on a plastic petri dish into a 15 mm x 15 mm square mesh with a strand-to-strand distance of 2 mm, and used for printability assessment. The cartridge and environment temperatures were not controlled, but room temperature was maintained at ≈ 20 °C and relative humidity at 60%. After printing, the samples were either left in air to dry, immersed in a 1x PBS solution, or, if containing TCEP, incubated in a 10^−2^ M NaIO_4_ solution for two hours. The scaffolds annealed with NaIO_4_ were washed three times with 1x PBS and finally immersed in 1 x PBS before further studies.

### 4.13. Printability assessment

A printability index (Pr), based on scaffold pore perimeter (L) and area (A), was used as a quantitative assessment of shape fidelity, which is defined as^30^:

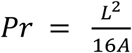

A Pr value of 1 indicates high geometric accuracy, corresponding to a square-shaped pore. Deviations from this ideal square geometry are reflected in Pr values less than 1 (indicating a more rounded shape) or greater than 1 (indicating an irregular shape). Optical images of printed constructs were analyzed using the ImageJ software to determine the pore perimeter and area, and consequently the Pr value (n=6).

### 4.14. Cryo-scanning electron microscopy (cryo-SEM)

Printed strands of granular hydrogels on glass were vitrified in undercooled liquid ethane using a Gatan (Pleasanton, OR, United States) CP3 plunge-freezer and transferred under liquid nitrogen to a home-made cryo-SEM sample holder^33^. The strand was cut with a blade cooled in liquid nitrogen, and the cross-section was visualized after tilting the sample. Secondary electron images were collected in high vacuum mode at 3 kV accelerating voltage under cryo-conditions using an FEI (Hillsboro, OR, United States) Quanta 400 FEG scanning electron microscope. Printed scaffolds seeded with HDF after 14 days of culture were fixed with 10 % formalin solution (Sigma-Aldrich, HT501128, Netherlands) for 15 min. Next, they were washed twice with 1x PBS and immersed in liquid nitrogen and transferred to a home-made cryo-SEM sample holder. Secondary electron images were collected in high vacuum mode at 3 kV accelerating voltage under cryo-conditions using an FEI (Hillsboro, OR, United States) Nova NanoSEM 450 scanning electron microscope.

### 4.15. Cell Culture

Primary the human dermal primary fibroblasts (HDFs) derived from adult skin (P10856, Innoprot, Spain) were cultured at a recommended seeding density of 5000 cells cm^−2^. DMEM (31966021, Gibco) supplemented with 10% fetal bovine serum (F9665, Sigma-Aldrich), 1% penicillin–streptomycin (10,000 U mL^−1^, 15140148, Gibco), and 1 ng mL^−1^ fibroblast growth factors (PHG0367, Gibco) was used as cell culture medium. HDF cells were cultured at 37 °C in a humidified atmosphere of 5% CO_2_. The medium was refreshed every 2 days until the cells reached over 90% confluency.

### 4.16. Cell Culture on Scaffolds

The dried microgels were sterilized via UV for 1 hour before being dispersed in 1x PBS, mixed, and jammed. For the indirect exposure, the ink was first printed in a single filament of 5mm length, crosslinked, and washed 3 times with 1 x PBS. Then, HDF cells at passages 6 and 8 were seeded at the density of 5,000 cells/cm^2^ (9,500 cells per well) in the 24-well plate (Sarstedt, 83.3922.500, the Netherlands). After one day in culture, the media was changed and the printed strands were added to the wells. Cell viability was checked on day 3, 7, and 14 with live/dead assay. 6 control samples per day were added: 3 positive – untreated control, and 3 negative: treated with ethanol.

For the direct exposure viability assay, 100 μL of the 20.0 wt.% granular hydrogel ink post-TCEP addition was cast on the bottom of a suspension 24-well plate, and the plate was centrifuged to obtain flat coverage and homogeneous spreading of the granular hydrogel (1500 rcf, 5 min). Afterwards, it was crosslinked with NaIO_4_ for two hours, and then washed 3 times with 1x PBS. The casts were incubated overnight at 37 °C with FBS, and HDF cells were seeded on top the day after. Cells were added at a volume of 100 µl to each scaffold and incubated for 60 minutes in the incubator for initial cell attachment, and after that, 1.5 mL of medium was added. After 14 days, live/dead assay was performed. Subsequently, a scaffold of the 20.0 wt.% ink with TCEP was 3D printed with 4 layers at dimensions 9 x 9 mm with 0.3 mm infill distance on 10 × 10 mm square coverslips and crosslinked with NaIO_4_ for two hours. After printing, the scaffolds were washed, incubated, and seeded as mentioned before for the casting of granular hydrogels. Live/dead and alamarBlue measurements of the metabolic activity were performed on day 3, 7, and 14. HDF cells were used between passages 6 and 8. To each printed scaffold a density of 8. 80,000 cells (42,105 cells/cm^2^) was use. For 2D controls (just HDF seeded in round glass coverslips (12 mm diameter; CB00120RA120MNZ0, Epredia) 9,500 cells (5,000 cells/cm^2^) without scaffolds) were added. The medium was refreshed every 1-2 days.

### 4.17. Cell viability

The cell viability on casted granular hydrogel scaffolds after 14 days or printed granular hydrogel scaffolds after 3, 7, and 14 days was investigated via live/dead assay. In short, a stock solution of FDA (ThermoFisher Scientific, F1303, the Netherlands) in acetone (concentration of 5 mg/mL) was prepared and further diluted to 20 μg/mL in 1x PBS. A solution of propidium iodide (PI) (ThermoFisher Scientific, P1304MP, the Netherlands) was prepared in 1x PBS (concentration of 20 μg/mL). The cells were incubated with 0.1 mL of FDA solution and 0.030 mL of PI solution in the 24-well plates for 10 minutes, followed by 3 washes with 1x PBS. The samples were then imaged with a fluorescence microscope with temperature control (Leica Stellaris 8) and counted. Cell viability was calculated with the use of the following equation: % viability = [(number of live cells)/(total number of cells)]×100.

### 4.18. Metabolic Activity

The metabolic activity was checked with the alamarBlue assay. Briefly, the alamarBlue reagent (ThermoFisher Scientific, DAL1025, the Netherlands) was mixed with cell culture medium in a 1:10 ratio and added to the cells. The samples were incubated at 37 °C for 4 hours, followed by supernatant collection and measurement of the fluorescence at excitation 560 nm and emission 590 nm (Tecan Spark M10 or BioTek Synergy H1 multimode microplate reader with Gen5 3.08 software). The percentage of alamarBlue reduction was calculated with the use of the following equation: % reduction = [(fluorescence intensity of the treated sample at 590 nm – fluorescence of the untreated control at 590 nm) / (fluorescence of 100% reduced alamarBlue at 590 nm (autoclaved for 15 min) – fluorescence of the untreated control at 590 nm)] × 100.

### 4.19. Statistical analysis

All the calculated results are reported as the mean value ± standard deviation. AlamarBlue statistical significance was analyzed using one-way ANOVA and a post hoc Tukey test with Origin8 software. Differences with p < 0.05 were considered significant. *p < 0.05, **p < 0.01, ***p < 0.001, ****p < 0.0001; ns – not significant.

## Supporting Information

This paper contains supplementary information.

## Acknowledgments

The authors acknowledge the European Union (ERC, JAM2PRINT, 101161716). Views and opinions expressed are, however, those of the author(s) only and do not necessarily reflect those of the European Union or the European Research Council. Neither the European Union nor the granting authority can be held responsible for them.

## Author contributions

The manuscript was written through the contributions of all authors. All authors have approved the final version of the manuscript.

