## Supplementary information for "Chemically-induced Jamming and 3D printing of Granular Hydrogels: Microgels as Reservoirs of Volume"

* corresponding authors

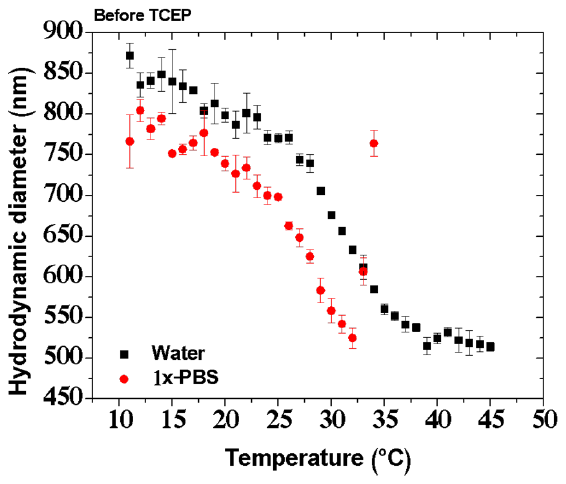

**Figure S1.** Evolution of the hydrodynamic diameter of the pristine microgels (before addition of tris-(2-carboxyethyl)phosphine hydrochloride, TCEP) as a function of the temperature in deionized water (black squares) and in 1x-PBS (red circles). The microgel concentration is 0.1 mg.mL^-1^ (0.01 wt.%). Above 32 °C in 1x-PBS, the microgels aggregate as the high salt concentration (around 0.137 M NaCl) screens the charges coming from the persulfate initiator (APS) on the surface of the particles. The colloidal stability above the Volume Phase Transition Temperature (VPTT) is then lost, and the particles aggregate.^34,35^

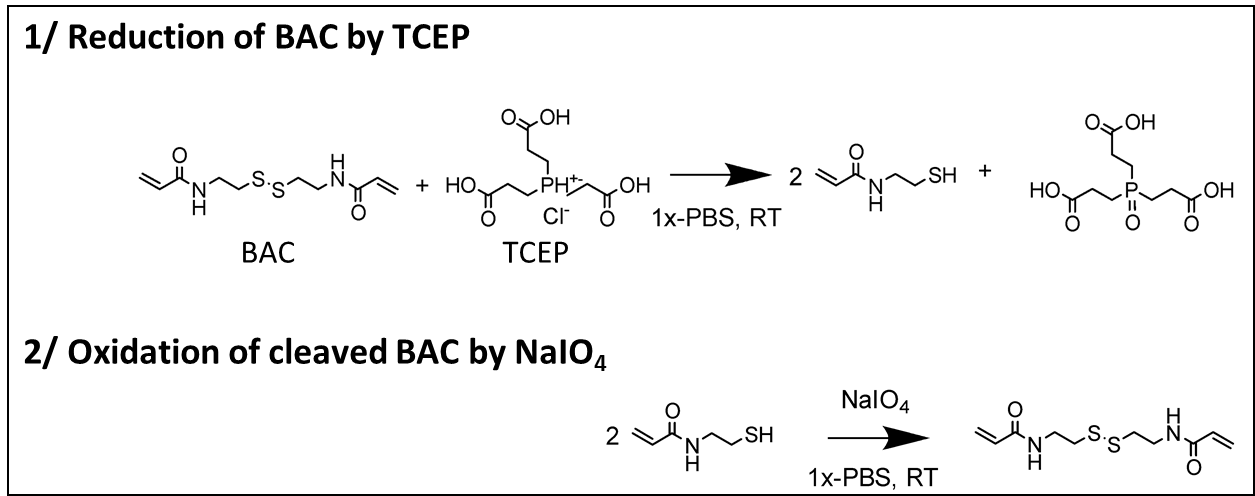

**Figure S2.** Reaction schemes of the reduction of the *N*,*N’*-bis(acryloyl)cystamine (BAC) by TCEP and of the oxidation of the resulting thiol groups by sodium (meta)periodate (NaIO_4)_.

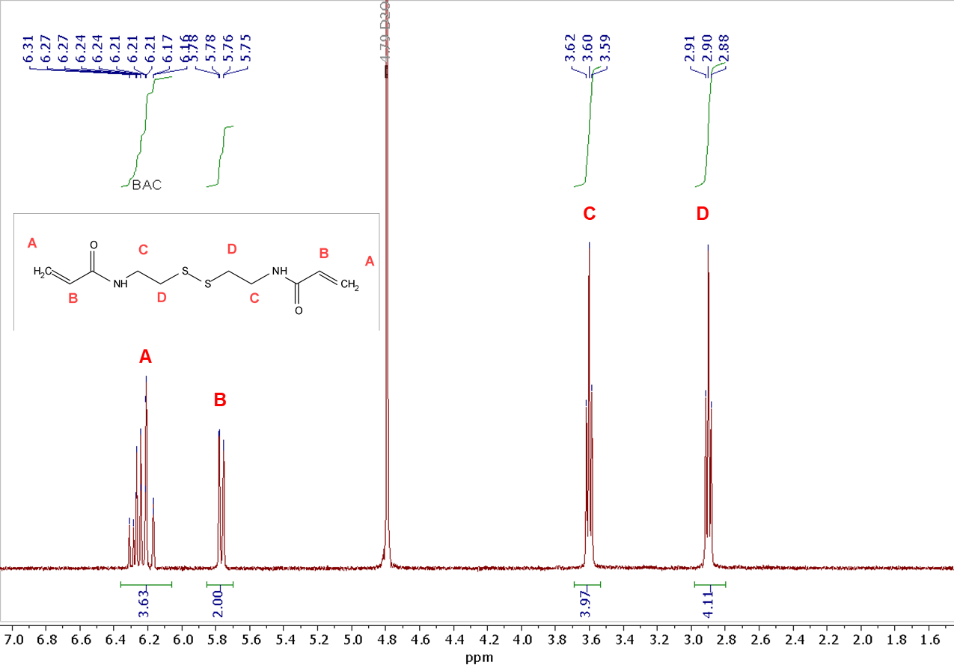

**Figure S3.** ^1^H NMR spectrum of BAC recorded in deuterated 1x-PBS.

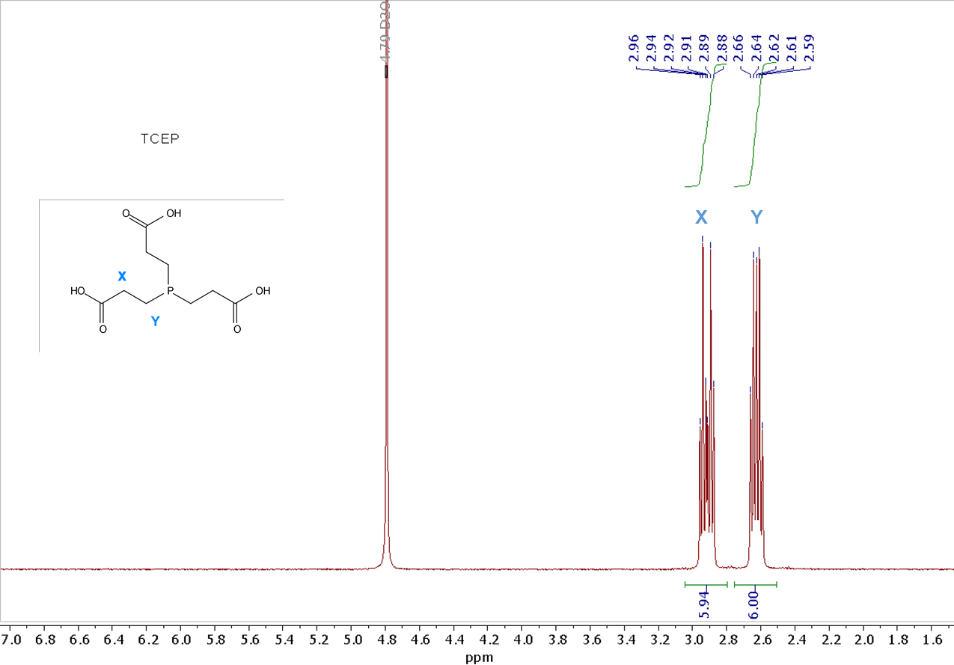

**Figure S4.** ^1^H NMR spectrum of TCEP recorded in deuterated 1x-PBS.

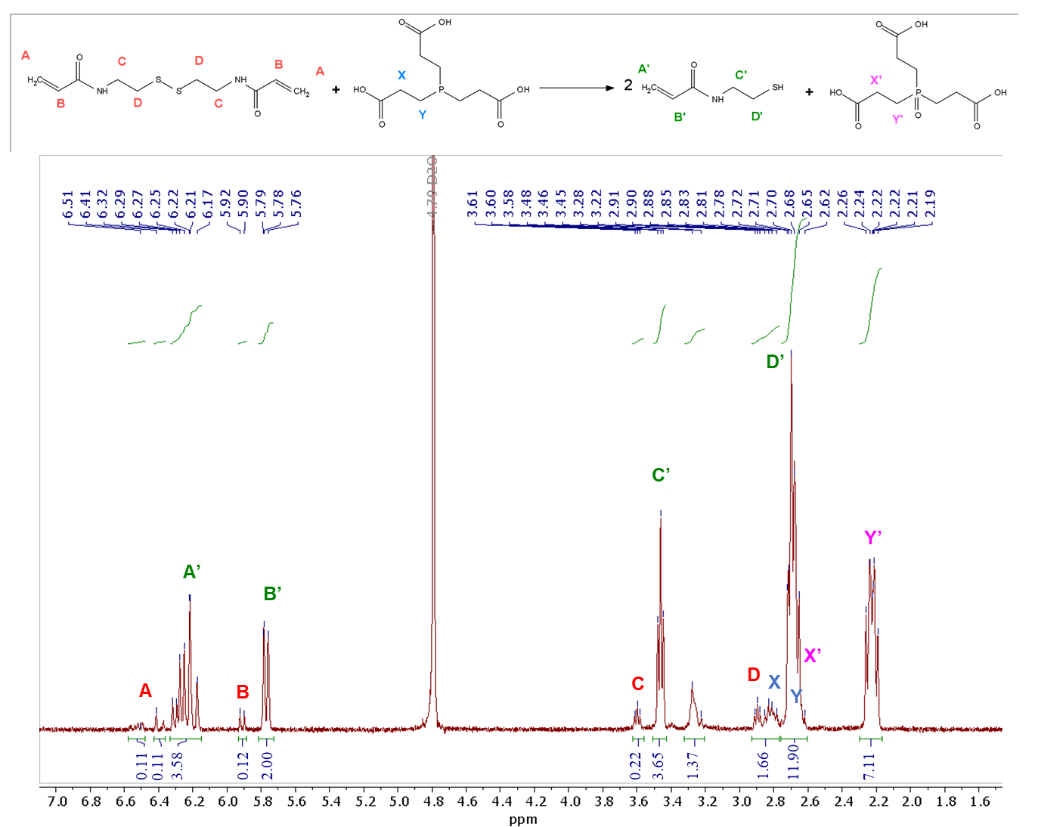

**Figure S5.** ^1^H NMR spectrum of 1:1 BAC:TCEP mol ratio 15 minutes after addition of TCEP recorded in deuterated 1x-PBS. The yield of the conversion was determined to be ≈ 95% after 15 minutes by calculating the ratio of the integrals of the peaks B’ and B that are respectively related to the α-protons of the acrylate moieties in BAC and in the product of the reduction of BAC by TCEP.

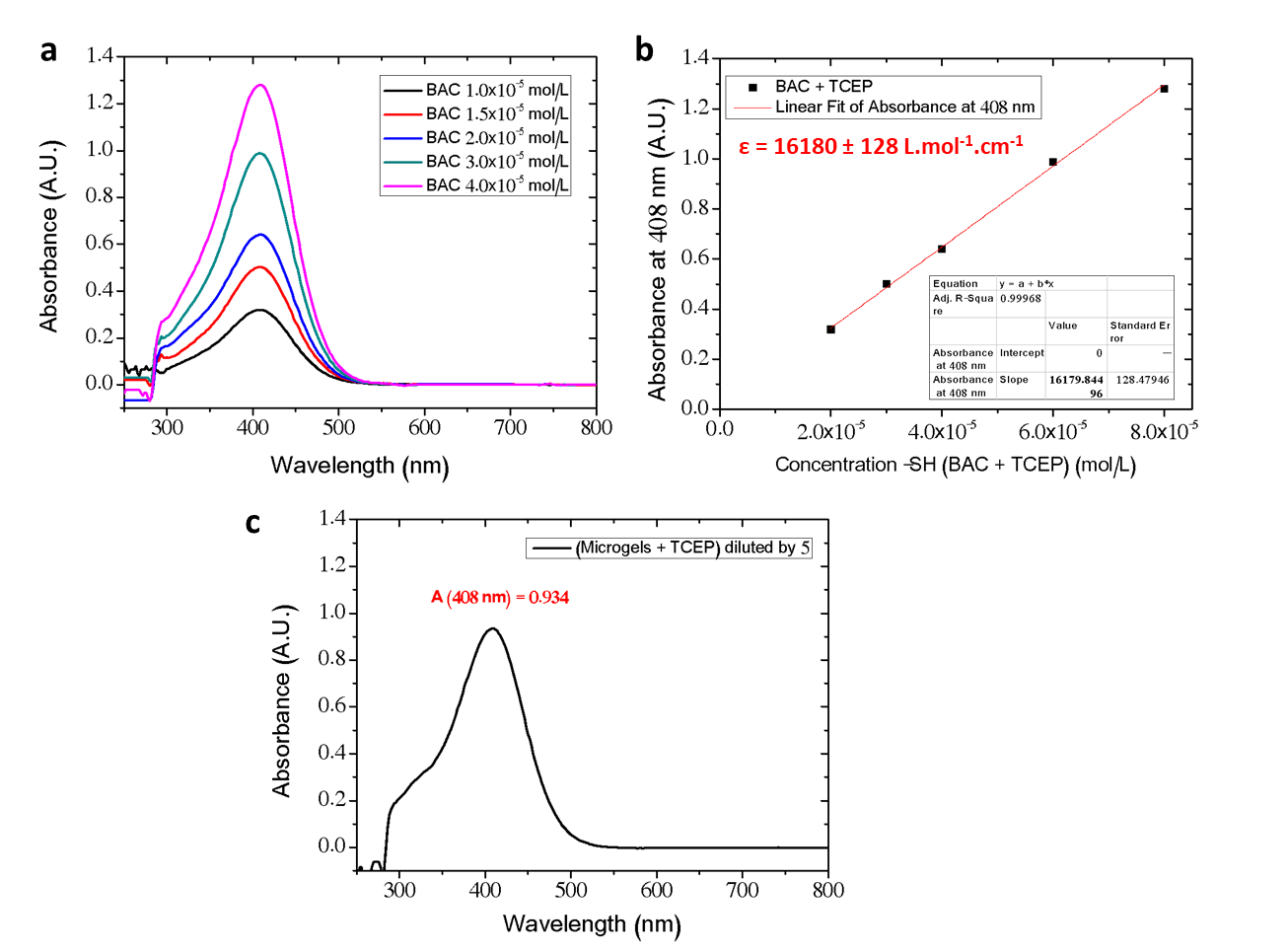

**Figure S6.** UV-visible characterization of BAC, TCEP and DTNB **(a)** UV-visible spectra of the (BAC + TCEP + DTNB) solution in 1x-PBS with BAC concentrations increasing from 1.0 10^-5^ M (black line) to 1.5 10^-5^ M (red line), to 2.0 10^-5^ M (blue line), to 3.0 10^-5^ M (green line), to 4.0 10^-5^ M (pink line). The concentration of TCEP and DTNB is 10^-3^ M (in excess compared to BAC). The absorption peak at 408 nm corresponds to the absorption of TNB produced by quantitative reaction with thiol groups obtained by the reduction of BAC by TCEP. **(b)** Derived calibration curve connecting the concentration of thiols to the absorption value of the peak at 408 nm. **(c)** UV-visible spectrum of the 5-times diluted supernatant of a (0.25 mg/mL microgel + 10^-3^ M TCEP + 10^-3^ M DTNB) dispersion in 1x PBS. The calculation of the BAC content in the microgels can be found in **Table S2**.

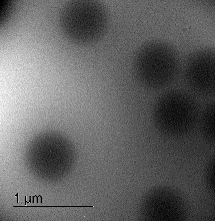

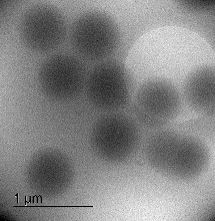

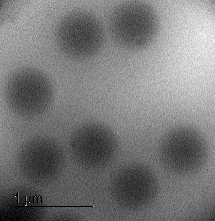

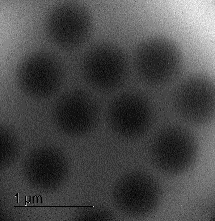

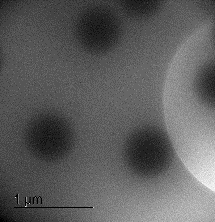

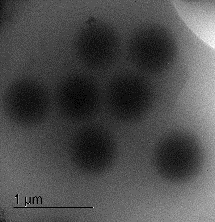

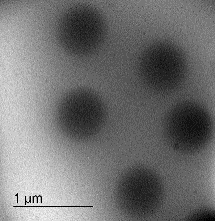

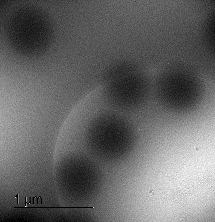

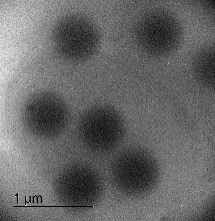

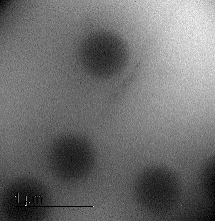

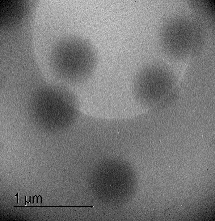

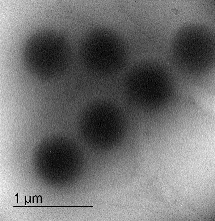

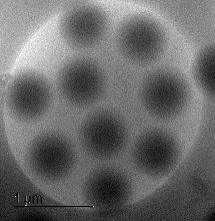

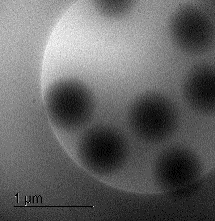

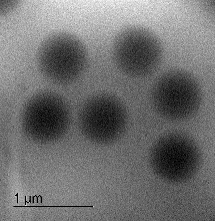

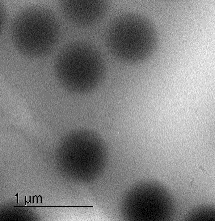

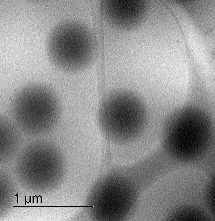

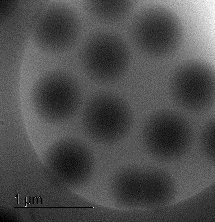

**a)**

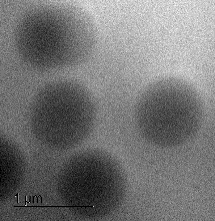

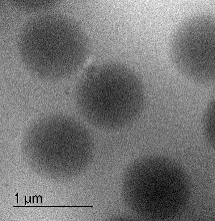

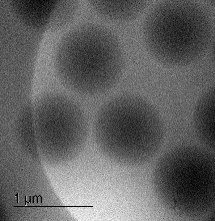

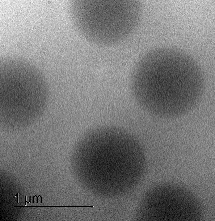

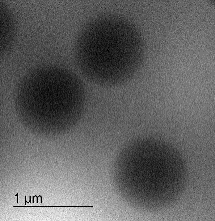

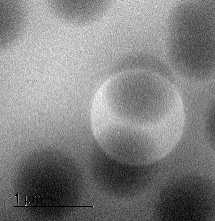

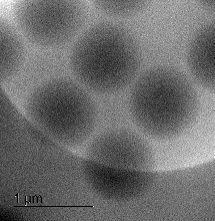

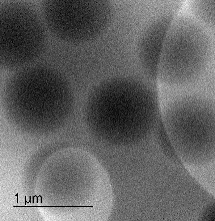

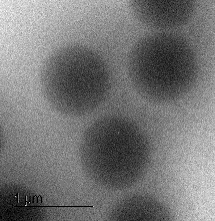

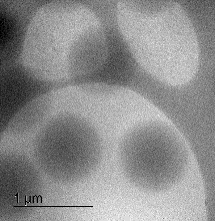

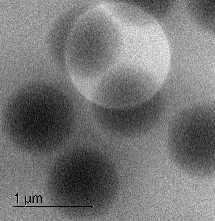

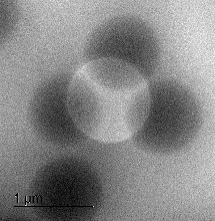

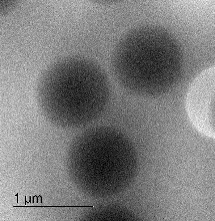

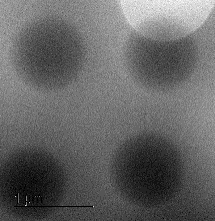

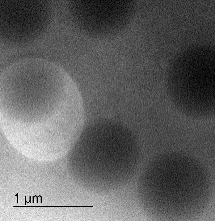

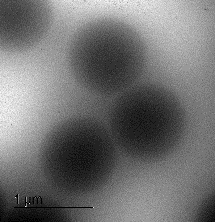

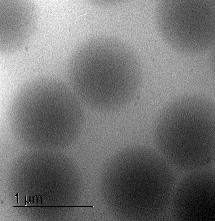

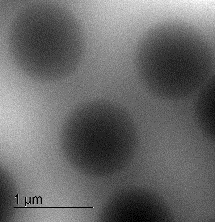

**b)**

**Figure S7.** Cryo-TEM images of dilute microgel dispersions **(a)** before and **(b)** after addition of TCEP. Scale bar is 1 μm.

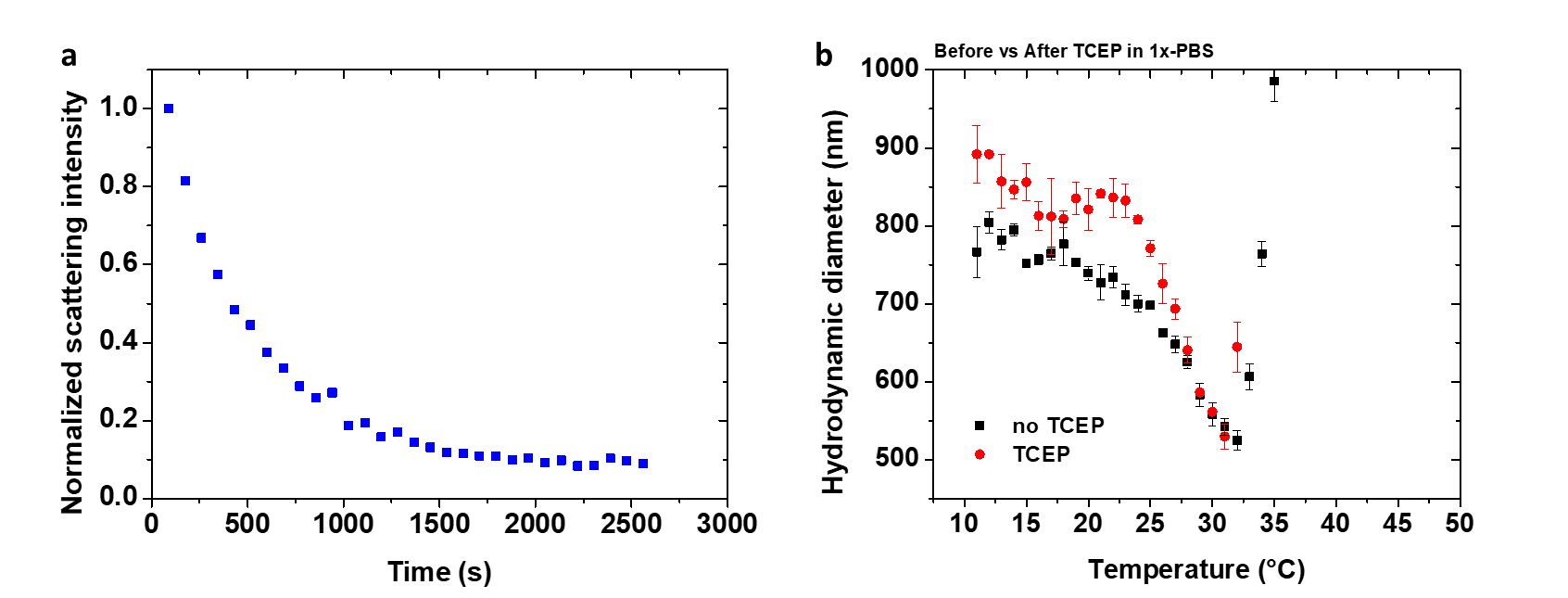

**Figure S8.** **(a)** Evolution of the scattering intensity (normalized by the scattering intensity at t = 0 s) as a function of time after addition of TCEP in 1x-PBS. **(b)** Evolution of the hydrodynamic diameter of the pristine microgels in 1x-PBS before addition of TCEP (black squares) and two hours after addition of TCEP (red circles) as a function of the temperature. The microgels concentration is 0.1 mg.mL^-1^ (0.01 wt.%) and TCEP concentration is 10^-2^ mol.L^-1^ (100 eq. compared to disulfide bonds).

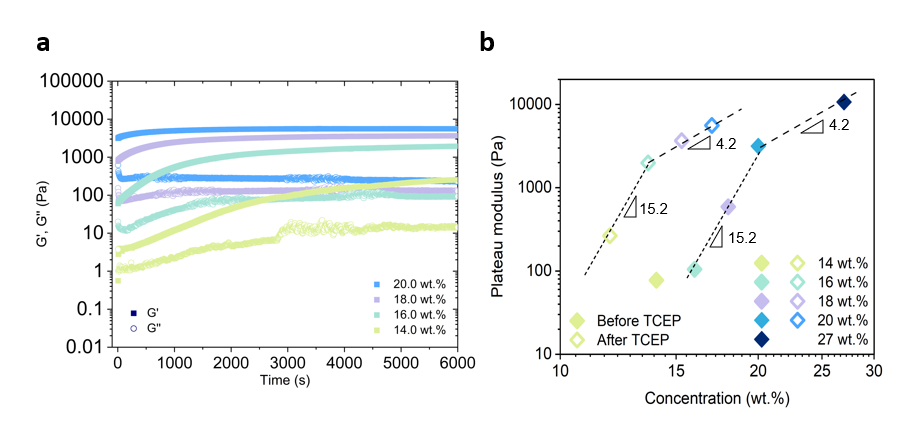

**Figure S9.** Rheological characterization of the microgel dispersions. **(a)** Time sweep (ω = 1 rad/s, γ = 1 % strain) upon jamming by TCEP addition to the microgel dispersions. **(b)** Plateau modulus G’ (determined at ω = 1 rad/s) as a function of the polymer concentration in the granular hydrogel. The dashed lines represent a linear fit of the dataset to the scaling prediction according to the theory of rubber elasticity. For the samples after TCEP reduction, the actual microgels concentration is rescaled by a factor 0.85 that takes into account the polymers chains release after chemical reduction. The higher concentrations regime (slope of 4.2) represents the macrogel-type behavior where the scaling of the hydrogel stiffness becomes independent of the granular nature, while the lower concentration regime (slope of 15.2) represents loosely packed network where granularity determines the stiffness.^36^ The granular hydrogels prepared at 16.0 wt.% and 18.0 wt.% are the only samples that transition from the loosely packed granular hydrogel behavior to the dense macrogel-type behavior after addition of TCEP. This may explain why these samples exhibit the highest relative increase of stiffness, yield stress, and consequently, printability upon chemical reduction.

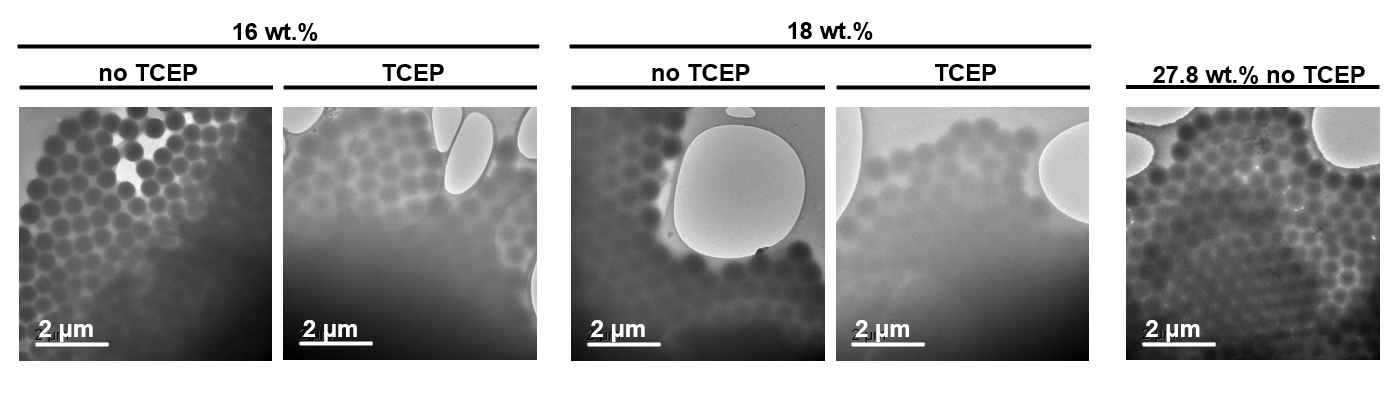

**Figure S10.** Cryo-TEM images of the 16.0 wt.%, 18.0 wt.%, and 27.8 wt.% granular hydrogels at the native state (no TCEP) and 6 000 s after addition of TCEP (TCEP).

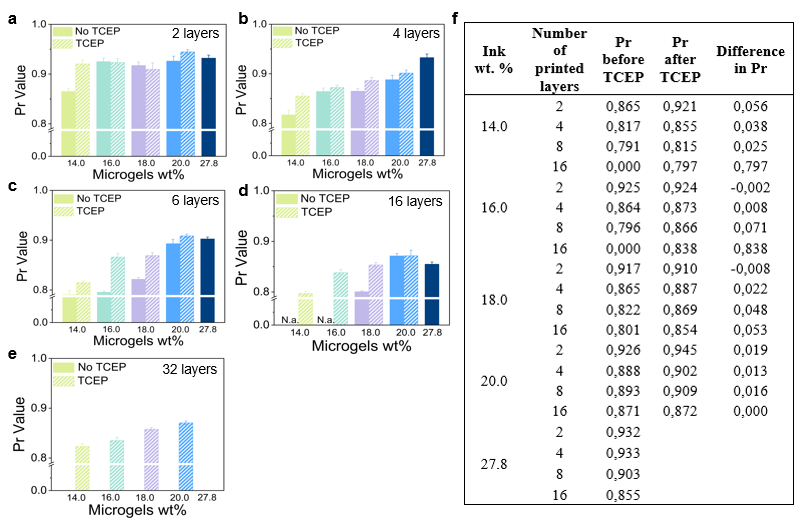

**Figure S11.** Printability indexes (Pr) calculated from Equation 1 (*Experimental Section*) for scaffolds printed using different microgel volume fractions, composed of 2 **(a)**, 4 **(b)**, 8 **(c)**, 16 **(d)**, and 32 layers. Pr differences before and after addition of TCEP **(f)**.

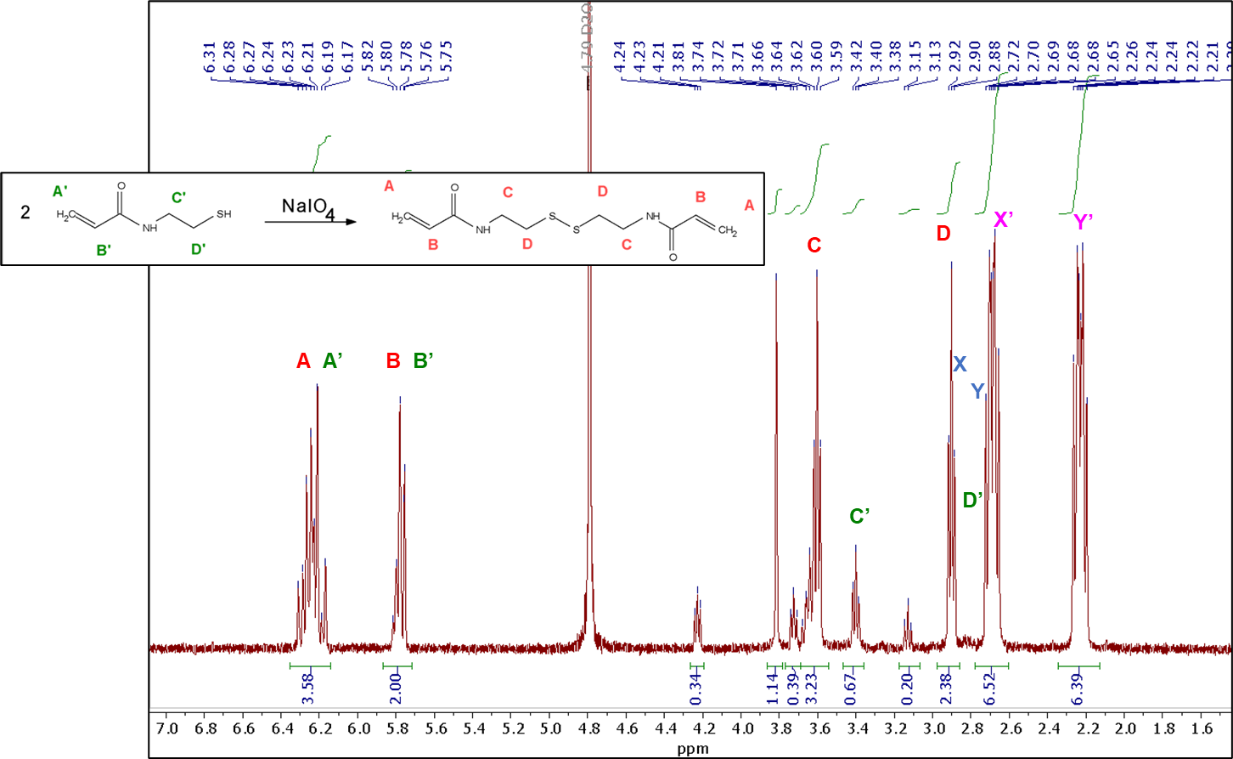

**Figure S12.** ^1^H NMR spectrum recorded in deuterated 1x-PBS of the original 1:1 BAC:TCEP mol ratio mixture 15 minutes after addition of 2 equivalents of NaIO_4_. The yield of the reaction was determined to be ≈ 82% after 15 minutes by calculating the ratio of the integrals of the peaks C and C’ that are respectively related to -CH_2_-NH- protons of the BAC and in the product of the reduction of BAC by TCEP.

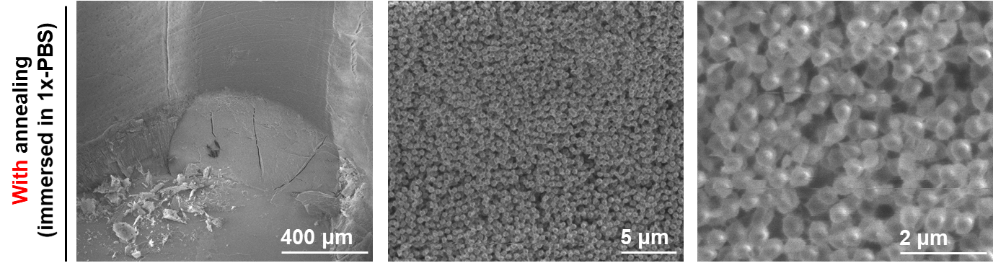

**Figure S13.** Cryo-SEM images (at different magnifications) of the cross-section of a 3D printed strand of the 20.0 wt.% with TCEP after annealing with NaIO_4_ and immersion in a 1x PBS solution.

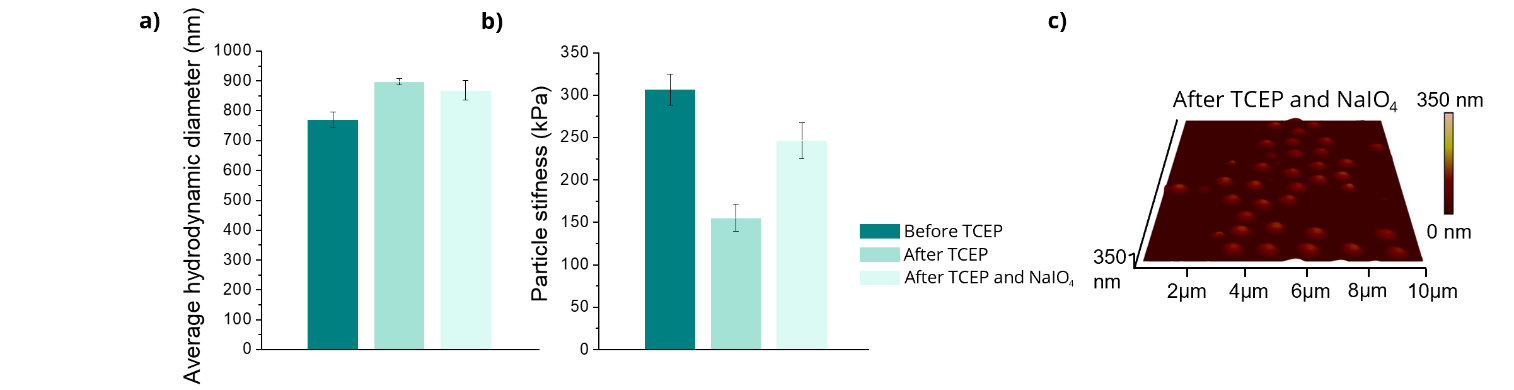

**Figure S14**. **(a)** Average hydrodynamic diameter and **(b)** stiffness of the microgels, before reduction with TCEP, two hours after TCEP addition, and after TCEP addition followed by the addition of NaIO_4_ for 5 minutes. **(c)** Wet-AFM images of the height of dilute microgel dispersions after chemical reduction with TCEP for two hours and oxidation with NaIO_4_ for 10 minutes.

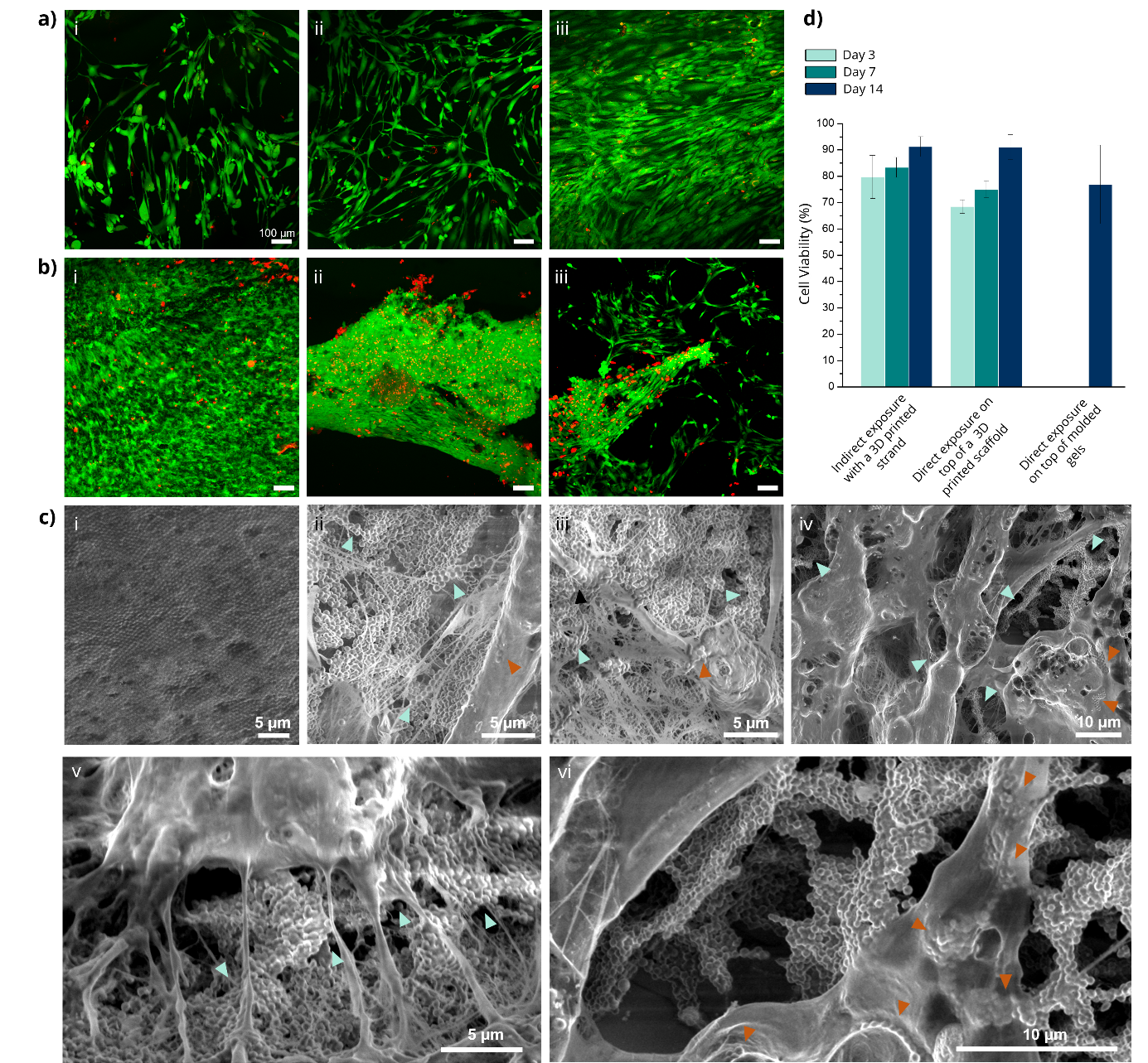

**Figure S15**. Cytocompatibility studies via indirect exposure **(a)** of HDF cells seeded on 24-well plate with a printed strand of pNiPAM granular hydrogel (20.0 wt. %), after (i) 3, (ii) 7, and (iii) 14 days of culture; and direct exposure **(b)** of HDF cells seeded on top of wells coated with the pNiPAM granular hydrogel (20.0 wt. %) after 14 days. **(c)** Cryo-SEM of a printed scaffold of pNiPAM granular hydrogel (20.0 wt. %) without (i) and with (ii-vi) HDF seeded on top after 14 days of incubation in media. (d) Cell viability calculations for HDF cells incubated with an annealed printed strand (indirect contact), on top of a printed scaffold (direct exposure print), and on top of molded granular hydrogels annealed (direct exposure mold).

**Table S1.** Compositions of the microgels dispersions used for rheology and 3D printing experiments.

| **Microgels concentration** | **Mass microgels (mg)** | **Mass 1x-PBS (mg)** | **Mass BAC (mg)** | **Mol BAC (mol)** | **Mass TCEP (mg)** |
| --- | --- | --- | --- | --- | --- |
| **27.8 wt.%** | 770.1 | 2000 | 96.6 | 3.7 10^-1^ | / |
| **20.0 wt.%** | 500.0 | 2000 | 62.7 | 2.4 10^-1^ | 69.0 |
| **18.0 wt.%** | 439.1 | 2000 | 55.08 | 2.1 10^-1^ | 60.6 |
| **16.0 wt.%** | 380.9 | 2000 | 47.8 | 1.8 10^-1^ | 52.6 |
| **14.0 wt.%** | 325.5 | 2000 | 40.9 | 1.6 10^-1^ | 45.0 |

**Table S2.** Data obtained from UV-Vis experiments presented in **Figure S6**. For the titration of BAC within the microgels, 2 mg of microgels were dispersed in 8 mL 1x-PBS (0.25 mg/mL).

| **A_408nm_ 5x-diluted (A.U.)** | **A_408nm_**  **(A.U.)** | **ε_TNB_**  **(L.mol^-1^.cm^-1^)** | **[-SH]**  **(mol.L^-1^)** | **[BAC]**  **(mol.L^-1^)** |
| --- | --- | --- | --- | --- |
| 0.94 | 4.67 | 16 180 | 2.89 10^-4^ | 1.45 10^-4^ |
| **BAC content**  **(mol)** | **Mass BAC**  **(mg)** | **NiPAM content**  **(mol)** | **mol%**  **BAC vs NiPAM** | **wt.% BAC** |
| 1.24 10^-6^ | 3.22 10^-1^ | 1.49 10^-5^ | **8.3 mol%** | **16.1 wt.%** |
